# Multivariate integration of histological images and gene expression data: a comparative review

**DOI:** 10.64898/2026.06.02.729734

**Authors:** Chengyi Ma, Jiadong Mao, Kim-Anh Lê Cao

## Abstract

Integrating histological images with gene expression data offers a promising approach for linking tissue morphologies to molecular signatures and improving disease subtyping. However, such integration remains challenging due to the high dimensionality of these datasets, cross-modal heterogeneity, and limited interpretability. Multivariate methods such as Sparse Canonical Correlation Analysis (Sparse CCA), Joint Nonnegative Matrix Factorisation (Joint NMF), and Angle-based Joint and Individual Variation Explained (AJIVE), have been used to address these challenges by reducing dimensionality while identifying features associated with latent factors, thereby enhancing biological interpretability. Despite increasing application in imaging-omics research, systematic comparisons of their methodological properties remain limited. Consequently, users often lack guidance on how to appropriately select these methods in practice, and these approaches are frequently treated as interchangeable despite differing modelling assumptions. Here, we use paired H&E images and gene expression data from breast cancer as a representative case study to examine the methodological characteristics, interpretability, and complementary properties of these integration approaches. Our results show that each method captures distinct yet complementary aspects of the underlying information. Although the biological findings are derived from the TCGA-BRCA datasets, the methodological insights identified here extend more broadly to imaging-omics integration studies. Overall, this comparative review highlights the strengths and limitations of each approach and outlines considerations for future methodological development.

## 1 Introduction

Histological imaging, such as haematoxylin and eosin (H&E) staining, is one of the most widely adopted assays in cancer diagnostics and research. H&E images provide rich morphological information, including cell size, cell shape and extracellular structure [1, 2]. They can be obtained at relatively low cost and are routinely incorporated into clinical workflows for tumour grading and subtype classification [1, 3, 4]. Meanwhile, gene expression profiles, such as PAM50 genes [5], quantify molecular activity at the transcriptomic level. These molecular information are critical for understanding gene regulatory programs and signalling pathways that underlie tumour development and progression. Molecular information has also been used for predicting therapeutic responses, and guiding personalised treatment [6–8]. Each of these modalities captures a distinct layer of breast cancer biology.

However, while H&E images capture tissue architecture and cellular morphology, they do not directly reveal the molecular characteristics of tumours. Establishing links between morphological patterns and molecular profiles provides a more comprehensive characterisation of tumour heterogeneity and improving disease stratification [9–11]. Bridging this gap therefore represents an important direction in translational cancer research.

Despite its promise, integrating histological images and gene expression data presents several methodological challenges. First, both imaging and gene expression data are typically high-dimensional, with the number of features far exceeding the number of samples. This high-dimensional setting renders many classical statistical approaches not applicable [12]. Second, histological images and gene expression data are inherently heterogeneous, representing distinct layers of biological system. Such heterogeneity complicates the integration across modalities [13]. Third, interpretability remains a key challenge[14]: effective integration should enable interpretable relationships between specific imaging patterns and gene signatures. For example, distinct morphological structures may correspond to coordinated up- and down-regulation of genes, revealing gene expression programs that underlie these imaging phenotypes.

Various integrative approaches have been used to address these challenges in multi-view data analysis [9]. Among them, multivariate integration methods represent a major set of techniques and can be categorised into three classes: canonical correlation-based approaches, joint factorisation-based approaches, and joint and individual variation decomposition approaches [15–19]. However, these methods have not been systematically compared within a unified analytical framework. Consequently, it remains unclear whether they capture consistent cross-modal signals or lead to differing biological insights. To address this gap, we conducted a representative case study integrating histological images and gene expression data in breast cancer, illustrating the methodological characteristics, interpretability, and complementary properties of these approaches. Breast cancer was chosen as it is among the most extensively studied cancers, and the abundance of publicly available data and prior studies provides a valuable reference for validating the biological relevance of identified image patterns and gene signatures.

In this review, we examine methods from the three methodological categories within a unified analytical framework. We first summarise their key characteristics and modelling assumptions. We then describe the TCGA-BRCA datasets and evaluation strategies, followed by method-specific evaluation and cross-method comparison to assess the performance of each approach. Finally, we discuss methodological implications and future directions for integrative analysis.

## 2 Multivariate integrative methods

In this section, we provide a brief overview of three families of multivariate integrative methods. We first introduce the matrix factorisation framework and define the key terminology used throughout the paper. We then describe the principles underlying each of the three method families, followed by a comparison of their main characteristics.

We selected Sparse CCA, Joint NMF, and AJIVE as representative methods for two main reasons. First, these methods have been successfully applied to the integration of histological images and gene expression data, demonstrating their relevance in imaging-omics research [15–19]. Second, we included only methods with publicly available implementations to enable reproducible evaluation and methodological comparison. Methods lacking accessible implementations were excluded.

### 2.1 Matrix factorisation framework and terminology

Multivariate integrative methods can be formulated within a structured matrix factorisation framework, which decomposes a data matrix *X* ∈ ℛ*^n^*^×^*^p^* into a low-dimensional score matrix *U* ∈ ℛ*^n^*^×^*^K^* and a loading matrix *V* ∈ ℛ*^p^*^×^*^K^*, such that *X* ≈ *UV* ^⊤^, where *n* denotes the number of samples, *p* denotes the number of features, and *K* represents the dimensionality of the latent space. Each component *k* is characterised by a sample score vector *u_k_*(the *k*-th column of *U*) and a corresponding feature loading vector *v_k_*(the *k*-th column of *V*). The loading vectors enable interpretation by linking specific features to latent components. When extended to multi-view settings, matrix factorisation models identify latent factors across different data views, thereby revealing cross-modal associations [12, 20, 21] (Figure 1).

**Figure 1.**
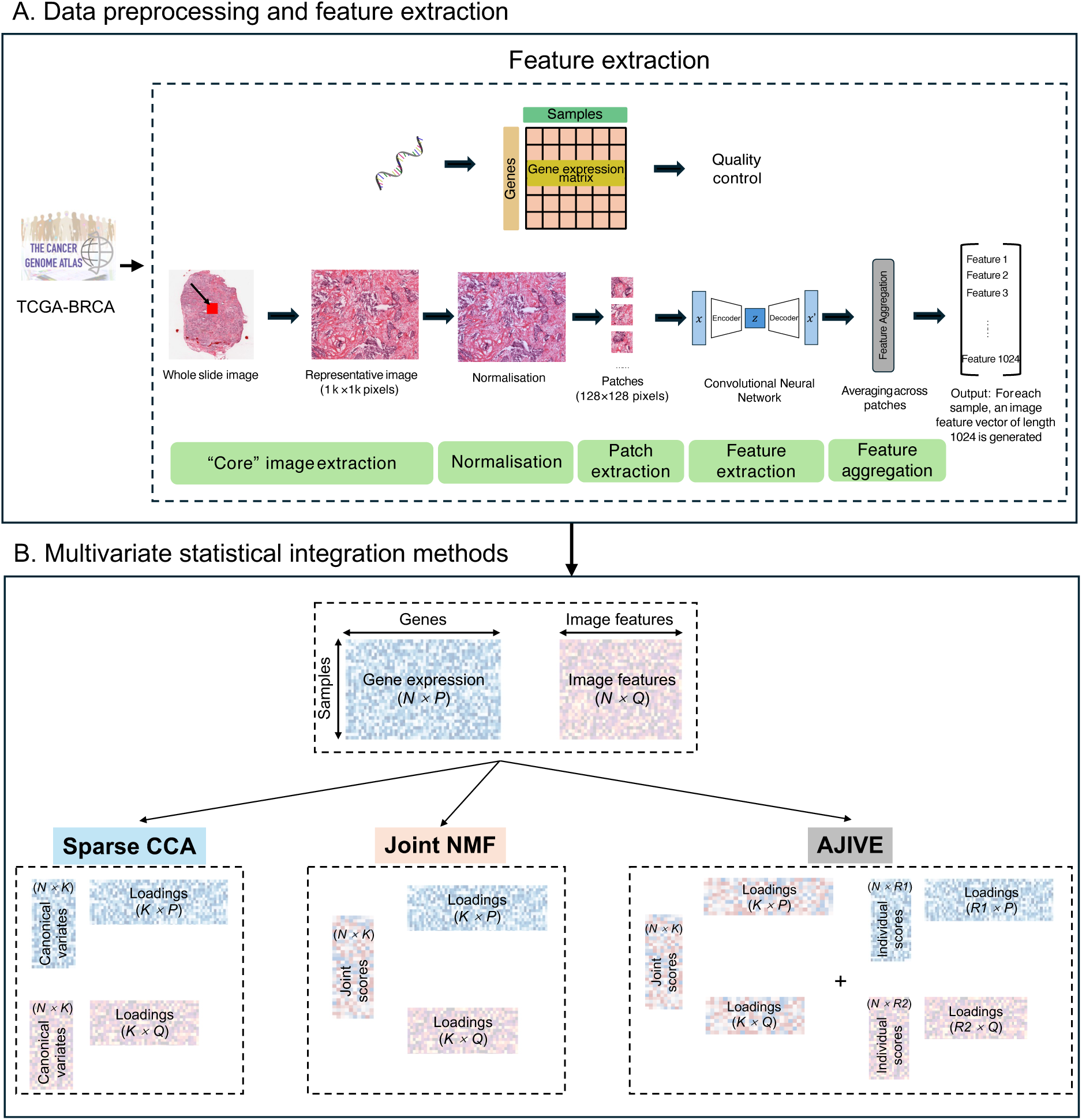
Workflow of image–gene expression data preprocessing and multivariate integration. **A:** Data preprocessing and feature extraction. Paired whole-slide images and gene expression were downloaded from TCGA-BRCA database. The study comprises 883 samples, each with a whole-slide image and paired FPKM-normalised gene expression profiles. Gene expression data were compiled into a data matrix and subjected to quality control. The image processing pipeline includes: (i) “core” image extraction, (ii) colour normalisation, (iii) patch extraction, (iv) feature extraction, and (v) feature aggregation. **B:** Multivariate statistical integration methods. Sparse CCA maximises the correlation between factors across modalities while imposing sparsity on the loading matrices. Joint NMF yields a joint score matrix together with loading matrices corresponding to each modality. AJIVE decomposes the data into both common and modality-specific structures, each represented by a corresponding score matrix and loading matrix.

To avoid ambiguity, we define the following terms used throughout the paper. The terms *block*, *data view*, and *modality* are used interchangeably to denote a data matrix corresponding to a specific data type measured on the same set of samples. A *component* (also referred to as a *latent factor* or *canonical variate*) represents an underlying latent factor identified by the method. The values of samples on a given component are referred to as the *score vector*, while the *loading vector* indicates the contribution of variables to the component. We use the term *multi-view* consistently to refer to data comprising multiple modalities. For example in this study, we focus on a two-view setting, consisting of histological image and gene expression data.

### 2.2 Canonical correlation-based analysis

Canonical Correlation Analysis (CCA) seeks to identify linear combinations of the original variables that maximise the correlation between two data views [22]. Specifically, given two data views *X*_1_ ∈ ℛ*^n^*^×^*^p^* and *X*_2_ ∈ ℛ*^n^*^×^*^q^* representing the gene expression matrix and the image feature matrix, respectively, where *p* and *q* denote the numbers of features in the two views, CCA identifies paired canonical variates across the two views, with canonical variates within each view being orthogonal to each other.

Let *u*_1_ and *u*_2_ denote the sample score vectors for *X*_1_ and *X*_2_, and *v*_1_ and *v*_2_ the corresponding loading vectors, CCA seeks *v*_1_ and *v*_2_ that maximise the correlation between the projected scores:

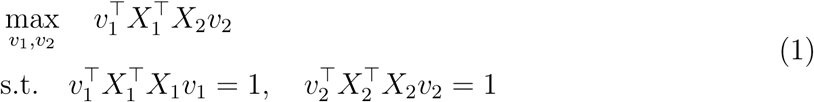

In high-dimensional settings, where *p, q* ≫ *n*, the cross-covariance matrix becomes singular, and traditional CCA is not applicable. To address this challenge, Sparse CCA [12] imposes sparsity on the loading vectors by adding penalties, shrinking the weights of non-relevant features to zero. Using the same formulation as in Equation (1), Sparse CCA solves:

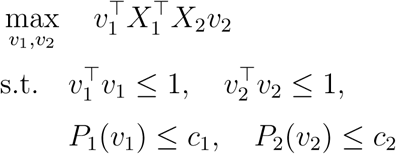

where *P*_1_ and *P*_2_ are convex penalty functions that induce sparsity in the loading vectors *v*_1_ and *v*_2_, respectively, with the *ℓ*_1_ penalty being the commonly used choice.

In practice, the feature covariance matrices 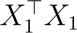 and 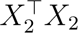 are typically not diagonal, resulting in latent score variances, 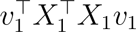 and 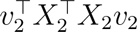 generally not equal to one. Consequently, Sparse CCA effectively maximises the covariance between the latent scores rather than their correlation. This behavior makes Sparse CCA more similar to Partial Least Squares (PLS), which seeks directions that maximise covariance between two data views [23].

To obtain multiple pairs of canonical variates, Sparse CCA adopts a deflation procedure. After extracting each pair of loading vectors (*v*_1_*, v*_2_), the cross-product matrix 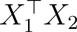 is updated by removing the rank-one approximation corresponding to the extracted component [24]. A process referred to as deflation [23]. This ensures that subsequent canonical variates capture new directions of association.

The sample score vectors are obtained as

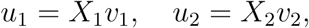

which can be interpreted as the projection of each sample onto the corresponding canonical variate.

From the perspective of the matrix factorisation framework introduced earlier, these projected score vectors correspond to the columns of the low-dimensional score matrices *U*_1_ and *U*_2_. Similarly, the columns of the low-dimensional loading matrices *V*_1_ and *V*_2_ correspond to the associated sparse loading vectors for each view. In contrast to classical CCA, which does not rely on deflation, Sparse CCA extracts components sequentially via a deflation procedure, making it more similar to PLS in practice.

### 2.3 Nonnegative matrix factorisation-based methods

Nonnegative Matrix Factorisation (NMF) constrains both the sample scores and feature loadings to be nonnegative. This constraint makes the *K* components interpretable and thus results in a parts-based representation [21]. The computation of NMF depends on three key components: error metric, initialisation, and search algorithm [25].

The error metric, also referred to as the objective function, is commonly defined by minimising the Frobenius norm between the original data and the reconstructed data. It is formulated as:

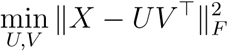

The initialisation step usually involves randomly initialising *U* and *V*, followed by iterative updates until convergence. However, the choice of initialisation can affect both the convergence speed and the final outcome, motivating ongoing efforts to develop more stable NMF algorithms.

Regarding search algorithms, common approaches include the multiplicative update method, alternating least squares, and gradient descent algorithms [21, 26–28]. They employ different optimisation strategies to improve convergence, making NMF adaptable to various data characteristics and application scenarios [29]. There is no universally accepted strategy for selecting the number of components *K*. In practice, incorporating prior knowledge about the data can help guide this choice [29].

Building upon traditional NMF, Zhang et al. [17, 30] have proposed Joint NMF, which simultaneously projected multiple views of data into a common space. Joint NMF ensures a shared joint score bases, while producing the corresponding loading matrices separately for each view. In the case of two data views, the objective function is formulated as follows:

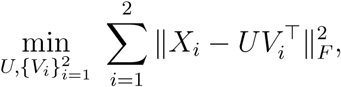

where each view 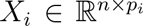 is approximated as 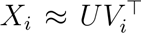. 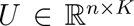 is the shared latent score matrix and 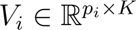 is the view-specific loading matrix.

### 2.4 Joint and individual variation decomposition approaches

Joint and Individual Variation Explained (JIVE) decomposes multi-view data into three components: shared (joint) signal, view-specific (individual) signal and noise [31]. This decomposition not only captures shared structures across multiple views but also reveals view-specific variation. Orthogonality constraints further ensure that joint and individual components are uniquely identified. The JIVE framework is summarised as follows.

Let 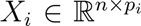 denote the *i*-th data matrix, where *n* is the number of samples and *p_i_* is the number of variables, with *i* = 1, 2. For example, *i* = 1 corresponds to the gene expression matrix, and *i* = 2 the image feature matrix. Each data matrix is decomposed as

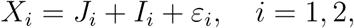

where *J_i_* represents the joint structure, *I_i_* the individual structure, and *ε_i_* the noise.

The joint and individual structures can be factorised as:

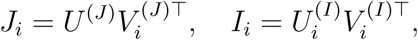

where *U* ^(*J*)^ ∈ ℛ*^n^*^×^*^K^* is the shared score matrix, and 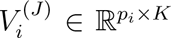 is the corresponding loading matrix for view *i*. Similarly, 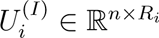 and 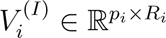 capture view-specific information, where *R_i_* is the individual rank for each view.

To ensure separation between shared and individual components, the following orthogonality constraint is imposed:

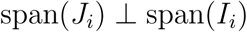

where span(·) denotes the subspace spanned by the columns of a matrix.

Feng et al. [32] developed angle-based JIVE (AJIVE) to improve the estimation of joint structure in multi-view data by using the Wedin bound and the random direction bound.

AJIVE has been shown to outperform JIVE on multi-view data, offering greater stability with heterogeneous data and improved computational efficiency [32]. The recently proposed DIVAS extension adopts a fully data-driven framework to identify partially shared structures, eliminating the need for prior rank specification [33, 34].

### 2.5 Comparison of methodological characteristics

To provide a structured comparison of the three integration methods, we summarise their key methodological characteristics in Table 1. The comparison includes objective functions, sparsity constraint, input requirements, outputs, interpretability, and computational properties.

**Table 1.** Comparative overview of Sparse CCA, Joint NMF, and AJIVE. *X*_1_ and *X*_2_ denote two views of data, with rows representing samples and columns representing features. *u*_1_ and *u*_2_ correspond to the latent factors for the two views in Sparse CCA, while *v*_1_ and *v*_2_ denote the associated loading vectors. For Joint NMF, *U* represents the shared joint score, and *V*_1_ and *V*_2_ correspond to the loading matrices for each view.

|  | Sparse CCA | Joint NMF | AJIVE |
| --- | --- | --- | --- |
| **Objective** | Maximise correlation between two views with sparsity | Joint low-rank non-negative factorisation across views | Decompose into joint (shared) and individual (view-specific) components |
| **Sparsity constraint** | Penalty on loadings | No | No |
| **Decomposition form** | Linear projections: $u_1 = X_1 v_1$; $u_2 = X_2 v_2$ | $X_1 \approx UV_1^\top$, $X_2 \approx UV_2^\top$, where $U$ is shared across views | $X = \text{Joint} + \text{Individual} + \text{Noise}$ |
| **Number of views** | Two (extendable via Sparse multi-CCA) | Extendable to multiple views | Natively supports multiple views |
| **Input requirements** | Paired data, mean-centred and scaled | Paired data, non-negative | Paired data, mean-centred and scaled |
| **Outputs** | $U_1, U_2$: Canonical variates<br>$V_1, V_2$: Sparse loadings | $U$: joint scores<br>$V_1, V_2$: loadings | $U^{(J)}$: Joint scores<br>$U_1^{(I)}, U_2^{(I)}$: Individual scores<br>$V_1^{(J)}, V_2^{(J)}$: Joint loadings<br>$V_1^{(I)}, V_2^{(I)}$: Individual loadings |
| **Interpretability** | Features can be selected via sparse loadings<br>Canonical variates capture maximally correlated patterns across views | Joint scores support sample clustering<br>Genes can be associated with corresponding factors | Joint scores capture common variation across views<br>Individual scores capture individual variation in each view |
| **Computational complexity** | Non-convex, alternating convex optimisation | Non-convex, alternating convex optimisation | Multiple SVDs and projections |

Sparse CCA, Joint NMF, and AJIVE differ fundamentally in their modelling objectives and the types of structures they capture. Sparse CCA is a correlation-driven approach that seeks directions maximising the association between paired views, while imposing sparsity on the loading matrices. Importantly, the method does not enforce a shared latent representation across data views, but instead identifies pairs of projections that are maximally associated. As a result, Sparse CCA typically produces paired latent factors with maximised positive associations between views.

In contrast, Joint NMF is a reconstruction-based method that performs joint low-rank nonnegative factorisation across data views with a shared latent factor matrix. Rather than directly maximising cross-view association, it aims to approximate the observed data, thereby enforcing a common latent space across views.

AJIVE adopts a subspace-based framework, explicitly decomposing each view into joint, individual, and noise components. Similar to Joint NMF, AJIVE enforces a shared representation by identifying a joint subspace common to all views. In contrast to Sparse CCA and Joint NMF, which primarily capture shared variation across data views, AJIVE explicitly decomposes the data into shared and view-specific components. The individual variation represents view-specific signals that are not shared by the other modality, and can be informative for identifying modality-specific patterns. In addition, AJIVE identifies shared low-dimensional subspaces rather than directly maximising cross-view correlation as in Sparse CCA. Therefore, the resulting joint components may exhibit either positive or negative associations between data views.

From a computational perspective, both Sparse CCA and Joint NMF involve non-convex optimisation and rely on iterative procedures. Sparse CCA uses regularisation parameters to control sparsity and is typically solved through alternating convex optimisation. Joint NMF employs multiplicative updates under nonnegativity constraints. In contrast, AJIVE is based on singular value decomposition and multiple projections to identify joint and individual subspaces.

Overall, the three methods differ in their objectives and decomposition forms. These differences have direct implications for the interpretability, particularly with respect to the biological relevance in cross-modal integration, which we examine in Section 5 and Section 6.

## 3 TCGA-BRCA data

This section describes the data source of this study and the preprocessing procedures (Figure 1A).

### 3.1 Data source

Paired bulk gene expression and H&E images from TCGA database (https://portal.gdc.cancer.gov/) were used for analysis. TCGA-BRCA data were downloaded using GDC Data Transfer Tool. Gene expression data were obtained by following query: Project=‘TCGA-BRCA’, Experimental Strategy=‘RNA-seq’, Data Category=‘transcriptome profiling’, Data Type=‘Gene expression Quantification’, Data Format=‘tsv’, Workflow

Type=‘STAR-Counts’. H&E images were obtained by following query: Project=‘TCGA-BRCA’, Experimental Strategy=‘Diagnostic Slide’, Data Category=‘biospecimen’, Data Type=‘Slide Image’, Data Format=‘svs’.

The first 10 characters of TCGA barcode (e.g., TCGA-XX-YYYY) represent the patient ID, which were used for screening paired samples across datasets. Paired gene expression data and whole-slide images (WSIs) for 883 samples were obtained.

### 3.2 Data preprocessing

#### Image preprocessing and feature extraction

The pipeline for WSIs included (i) 1000×1000 pixel image extraction, (ii) colour normalisation, (iii) patch extraction, (iv) feature extraction, and (v) feature aggregation (Figure 1).

#### 1000×1000 pixel image extraction

The extraction of 1000×1000 pixel images from WSIs was performed using ImageJ (https://imagej.net), following the approach described in [15]. The original H&E WSIs were in SVS format and were imported into ImageJ using the Bio-Formats plugin. A 1000×1000 pixel tile was selected if the mean grayscale value of both the tile itself and its surrounding tiles was below (darker) 190 out of 255.

#### Color normalisation of 1000×1000 pixel images

Color normalisation was applied to standardise the stain intensity of the extracted 1000×1000 pixel images [35]. Various factors can influence the intensity of H&E staining, such as the amount of dye used, different staining protocols, and difference in scanners. Without normalisation, these 1000×1000 pixel images exhibit noticeable visual color differences, which could potentially impact the accuracy of the extracted image features.

#### Training for convolutional autoencoder (CAE)

We built the CAE following the approach in [15]. The CAE is a model based on convolutional neural networks, which is commonly used in unsupervised learning. It consists of two main components: the encoder and the decoder. The encoder comprises 5 Conv2D layers followed by 5 MaxPooling2D layers, which progressively compress the input image into a compact latent representation. The decoder consists of 5 UpSampling2D layers and 5 Conv2D layers, which reconstruct the latent representation back into the original image. The network is trained with the objective of making the reconstructed image as close as possible to the input. The latent representation between the encoder and decoder encapsulates the core features of the image, where it contains 1024 features in this study.

Specifically, for each 1000×1000 pixel image, we applied random cropping and rotation for image augmentation, resulting in 128×128 pixel images. These images were first converted into array format before being fed into the CAE as input. In total, we trained the model for 3000 epochs.

#### Image feature extraction

We first loaded the fully trained weights into the CAE model. For each 1000×1000 pixel image, we randomly sampled 100 128×128 pixel images. We extracted the latent features using trained CAE and aggregated image features by averaging for each 1000×1000 pixel image. This procedure resulted in a feature matrix of 883 samples with 1024 features. The features with values greater than 0.001 in at least 20% of samples were retained, resulting in an image feature matrix of 883 samples by 563 image features for subsequent analysis. Examination of image feature density across samples revealed consistent patterns (Supplementary Figure S1).

#### Gene expression data preprocessing

Gene expression levels were quantified as Fragments Per Kilobase Million (FPKM) values. Genes with FPKM greater than 1 in at least 20% of samples were retained, and the 5000 most variable genes were subsequently selected. This procedure resulted in a gene expression matrix of 883 samples by 5000 genes for subsequent analysis. Examination of gene expression density across samples revealed consistent patterns (Supplementary Figure S1).

## 4 Evaluation strategies and biological interpretation

In this section, we outline the evaluation strategy for assessing the integration methods and their biological relevance. This includes quantifying the variance explained by each method, examining the consistency of selected genes across methods, and performing gene set enrichment and PAM50 molecular subtyping analyses.

### Variance explained by each method

We quantified the contribution of each factor to the original data. The scaled data view was projected onto the corresponding loading vector associated with a given factor. The squared norm of the resulting projection was then divided by the total variance of the scaled data view to estimate the proportion of variance explained. This procedure was applied consistently across all methods to allow a comparable assessment of factor contributions.

### Gene overlap across integration methods

For Sparse CCA and AJIVE, genes were first separated into positive and negative groups according to the sign of their loadings for each factor. Within each group, the top genes were selected based on the magnitude of their loading values. The selected genes were then combined to form the gene set for that factor. In Joint NMF, as all loadings are non-negative, the top genes for each factor were selected directly based on the magnitude of their loading values.

The gene sets associated with each factor from each method were then organised into a list, with each element representing the genes selected for a specific factor. We subsequently used the UpSetR package (v1.4.0) to visualise the intersections among these gene sets and to examine overlaps across different factors.

### Gene enrichment analysis

Gene sets for each factor were defined as described above. Gene Ontology (GO) analysis was then performed separately for the selected genes using the clusterProfiler R package (v4.14.6).

### PAM50 molecular subtyping

PAM50 molecular subtypes were predicted using the genefu R package (v2.40.0). Samples were classified into one of five intrinsic subtypes: Basal, HER2-enriched, Luminal A (LumA), Luminal B (LumB), and Normal-like. In the low-dimensional representations identified by each integration method, samples were coloured according to their PAM50 subtype to highlight subtype-specific structure. In addition, the magnitudes of loadings corresponding to PAM50 genes were examined across different methods.

## 5 Method-specific evaluation of image patterns and associated gene signatures

Following the construction of the gene expression and image feature matrices, we applied Sparse CCA, Joint NMF, and AJIVE to integrate the two views (Figure 1B). In this section, we evaluate the performance of each method in capturing associations between the two views on the TCGA-BRCA study.

### 5.1 Sparse CCA linked nuclear morphology to distinct molecular pathways

We examined the samples exhibiting the extreme positive or negative values along each canonical variate (Figure 2). This visual inspection provides insight into the information captured by each latent factor at the sample level. Moreover, since CCA factor pairs corresponding to the two views of data are positively correlated, samples with positive canonical variate values can be associated with the genes with positive loadings, particularly those with larger magnitudes. This, in turn, facilitates interpretation of the relationship between images and gene signatures.

**Figure 2.**
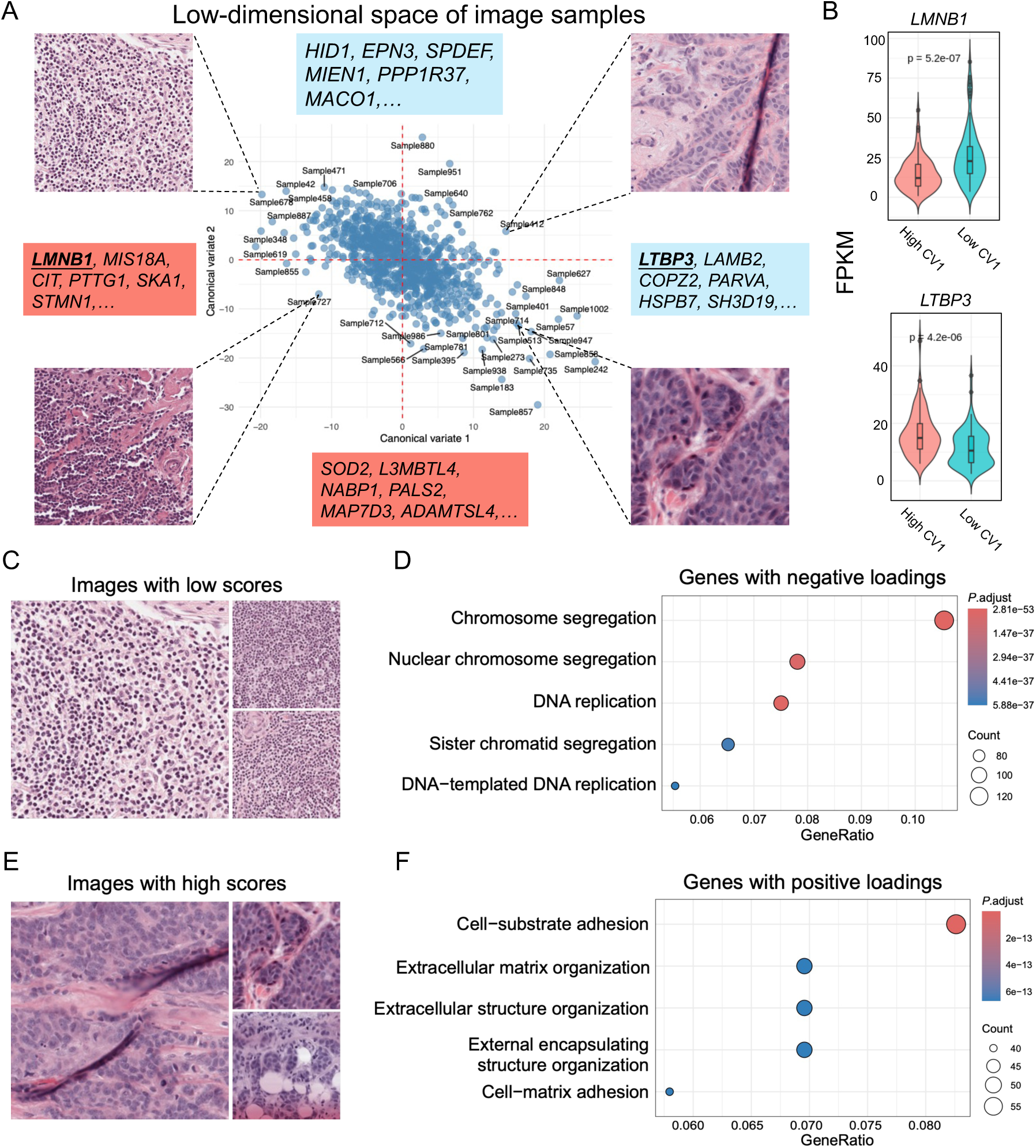
Joint canonical representation of H&E images and gene expression via Sparse CCA, with biological interpretability at the image and gene levels. **A:** Projection of the samples for the first two image canonical variates, with representative image shown for each quadrant. Top genes associated with either positive (blue) or negative (salmon) canonical variates corresponding to positive or negative loadings. **B:** Expression levels of two representative genes (*LMNB*1, *LTBP* 3) in subgroups defined by the top and bottom 10% of image canonical variate 1 values (CV1), showing significant differences. **C:** Representative images with extreme negative values of canonical variate 1 (low scores). The values of the samples projected onto a given canonical variate are referred to as the *scores*. **D:** Top five Gene Ontology terms enriched among genes associated with images with low scores (CV1). **E:** Representative images with extreme positive values of canonical variate 1 (high scores). **F:** Top five Gene Ontology terms enriched among genes associated with images with high scores (CV1). Sparse CCA identified a canonical variate contrasting small densely packed nuclei with larger nuclei, linked to distinct molecular pathways including chromosome segregation and DNA replication, or extracellular matrix organisation and cell-substrate adhesion.

We first examined two canonical variates for each data view (Supplementary Methods) and visualised the image samples in the resulting two-dimensional canonical variate space. Samples projected within each quadrant revealed that neighbouring images exhibited similar morphological patterns (Figure 2A, C and E), whereas samples from different quadrants displayed distinct characteristics (Figure 2A). Along the first canonical variate axis, samples with extreme negative score values displayed nuclei that were small and densely packed, whereas samples with extreme positive score values exhibited larger nuclei. Along the second canonical variate axis, samples with extreme negative score values were characterised by closely packed cells, with deeply stained (hyperchromatic) nuclei and a low cytoplasmic proportion. In contrast, those with extreme positive score values exhibited more loosely organised tissue, with noticeable gaps between cells, a higher proportion of stroma, and a relatively brighter tissue background (Figure 2A).

To investigate the relationship between image morphology and gene expression, we selected gene sets associated with each canonical variate. Genes with positive loadings corresponded to samples with positive scores on the same variate, whereas genes with negative loadings corresponded to samples with negative scores. For instance, image samples aligned with the negative direction of the first canonical variate were associated with genes such as *LMNB1*, *MIS18A*, *CIT*, *PTTG1*, *SKA1*, and *STMN1*, while those aligned with the positive direction were linked to *LTBP3*, *LAMB2*, *COPZ2*, *PARVA*, *HSPB7*, and *SH3D19* (Figure 2A). Notably, *LMNB1* encodes protein which plays a important role in maintaining chromatin structure [36]. Recent study have identified LMNB1 as a therapeutic target for breast cancer [37]. *LTBP3* encodes a protein essential for regulating TGF-*β* signaling and has been implicated in early metastatic events during cancer progression [38]. Similarly, images in the negative direction of the second canonical variate were associated with *SOD2*, *L3MBTL4*, *NABP1*, *PALS2*, *MAP7D3*, and *ADAMTSL4*, whereas image in the positive direction was linked to *HID1*, *EPN3*, *SPDEF*, *MIEN1*, *PPP1R37*, and *MACO1* (Figure 2A). SOD2 has been reported to be associated with aggressive subtypes of breast cancer and increased SOD2 level correlates with poorer outcomes [39, 40].

To investigate gene expression patterns across subgroups defined by each image canonical variate, samples were stratified according to their corresponding image score values. The score distributions for canonical variates 1 and 2 were approximately normal and centred around zero (Supplementary Figure S3). Based on this distribution, the top and bottom 10% of samples (i.e. those with the most extreme positive and negative scores) were selected and assigned to two subgroups for downstream analysis. We observed that the gene expression patterns differed markedly between these subgroups. For instance, in the first canonical variate, *LMNB1* expression was substantially higher in samples with negative scores (*p* = 5.2e-07, *t* -test), whereas *LTBP3* expression was elevated in samples with positive scores (*p* = 4.2e-06, *t* -test) (Figure 2B).

To further explore the functional relevance of the genes identified, we performed Gene Ontology (GO) term enrichment analysis on the genes associated with two group of images (images with low scores or high scores), respectively. Along the first canonical variate, genes corresponding to the images with low scores, characterised by small and densely packed nuclei (Figure 2C), were significantly enriched for biological processes such as chromosome segregation and DNA replication (Figure 2D). In contrast, genes associated with the high score images, characterised by larger nuclei (Figure 2E), were predominantly enriched for terms related to cell-substrate adhesion and extracellular matrix organisation (Figure 2F). These two subgroups of samples not only displayed distinct morphological features but were also associated with distinct gene signatures and enriched ontology terms. Collectively, these results indicate underlying biological differences between the sample groups defined by canonical variates.

### 5.2 Joint NMF identified two clusters with distinct glandular structures and gene signatures

We examined the two-dimensional shared joint score matrix (Supplementary Methods), which captures the shared information between the two data views. Clustering of the joint score matrix identified two clusters with maximum silhouette score (Supplementary Figure S4). We then examined the factor scores within each cluster. Overall, Cluster 1 displayed higher values for Factor 1, whereas Cluster 2 exhibited higher values for Factor 2 (Figure 3A).

**Figure 3.**
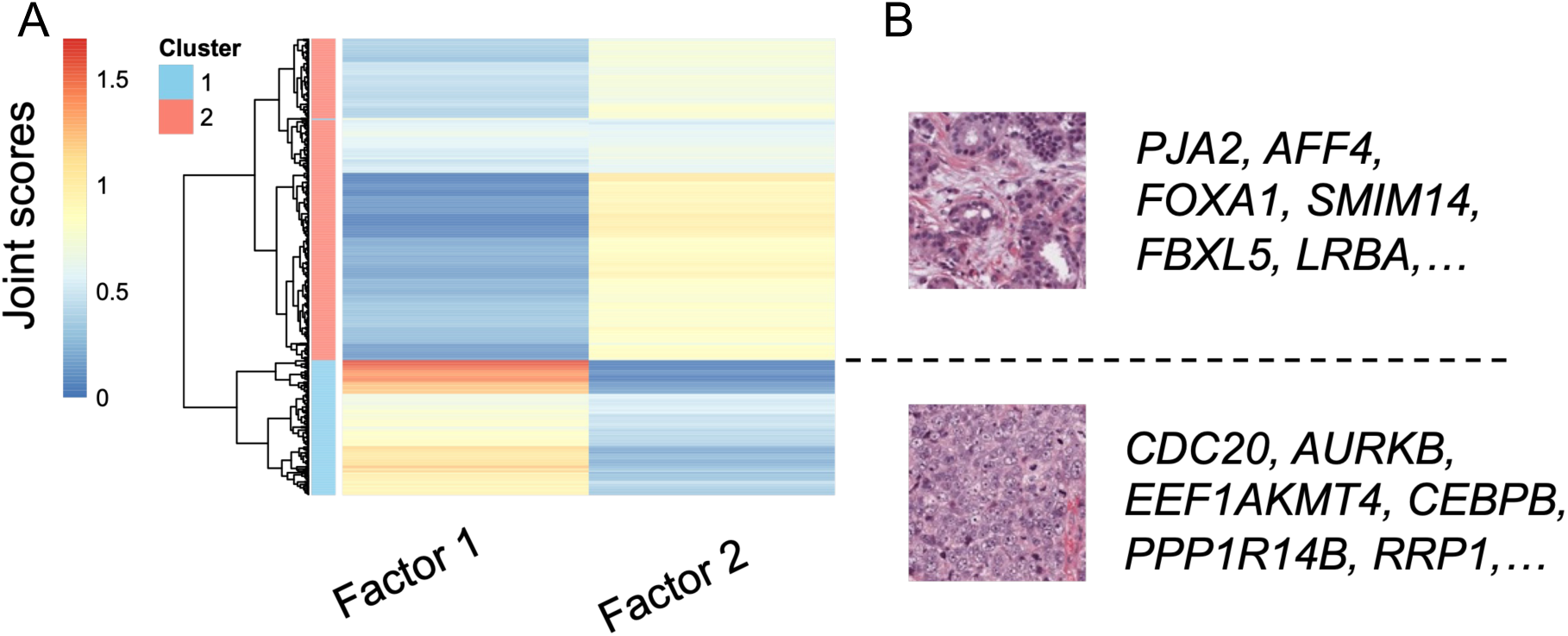
Latent factors derived from Joint NMF link H&E image patterns with gene sets. **A:** Heatmap of the joint scores obtained from Joint NMF. Cluster labels were assigned based on *k*-means clustering of the joint scores, revealing two stable sample clusters. **B:** Representative images and genes associated with each cluster. Joint NMF identified two clusters from the shared score matrix, with one cluster characterised by preserved glandular structures and stromal presence, and another showing nuclear atypia and loss of glandular architecture, together with distinct associated gene signatures.

To further understand the information captured by the factors, we compared representative images from each cluster. In Cluster 2, glandular structures were discernible, with some exhibiting tubular arrangements. Additionally, stroma was visibly present between the glandular structures. In contrast, Cluster 1 exhibited marked nuclear atypia, loss of glandular architecture, and less visible stroma, indicative of a higher-grade breast cancer (Figure 3B).

The gene loading matrix allowed us to link specific genes to individual factors. For each factor we extracted the genes with extreme positive loading values (all values are positive in Joint NMF). Factor 1 was associated with genes such as *CDC20*, *AURKB*, *EEF1AKMT4*, *CEBPB*, *PPP1R14B*, and *RRP1*, whereas Factor 2 was closely linked to *PJA2*, *AFF4*, *FOXA1*, *SMIM14*, *FBXL5*, and *LRBA* (Figure 3B). Notably, CDC20 expression has been reported to be elevated in tumor tissues from breast cancer patients and is associated with poor prognosis and reduced overall survival [41]. PJA2-associated immune microenvironment plays an important role in the progression of invasive micropapillary carcinoma (IMPC) of the breast [42]. AURKB plays a critical role in cell division, and targeting AURKB has been explored as a potential therapeutic strategy in triple-negative breast cancer (TNBC) [43].

### 5.3 AJIVE separated joint and individual structures, with joint components capturing nuclear morphology

We identified two joint components (Supplementary Methods and Supplementary Figure S5). In the first joint component, the paired joint scores from gene expression and image features were strongly negatively correlated (Supplementary Figure S6). This observation contrasts with Sparse CCA, where the paired canonical factors were always positively correlated (Supplementary Figure S2A and S2B). This difference likely reflects distinct modelling assumptions and optimisation strategies between the methods. A more detailed comparison of the similarities and differences across methods is provided in the next Section. We further examined image samples with either extreme positive or negative scores on this joint component. Image samples with extreme positive scores exhibited densely packed small nuclei, whereas image samples with negative scores displayed enlarged and heterogeneous nuclei, surrounded by small eosinophilic dots, with a large proportion of empty background visible in the field of view (Figure 4; Supplementary Figure S6).

**Figure 4.**
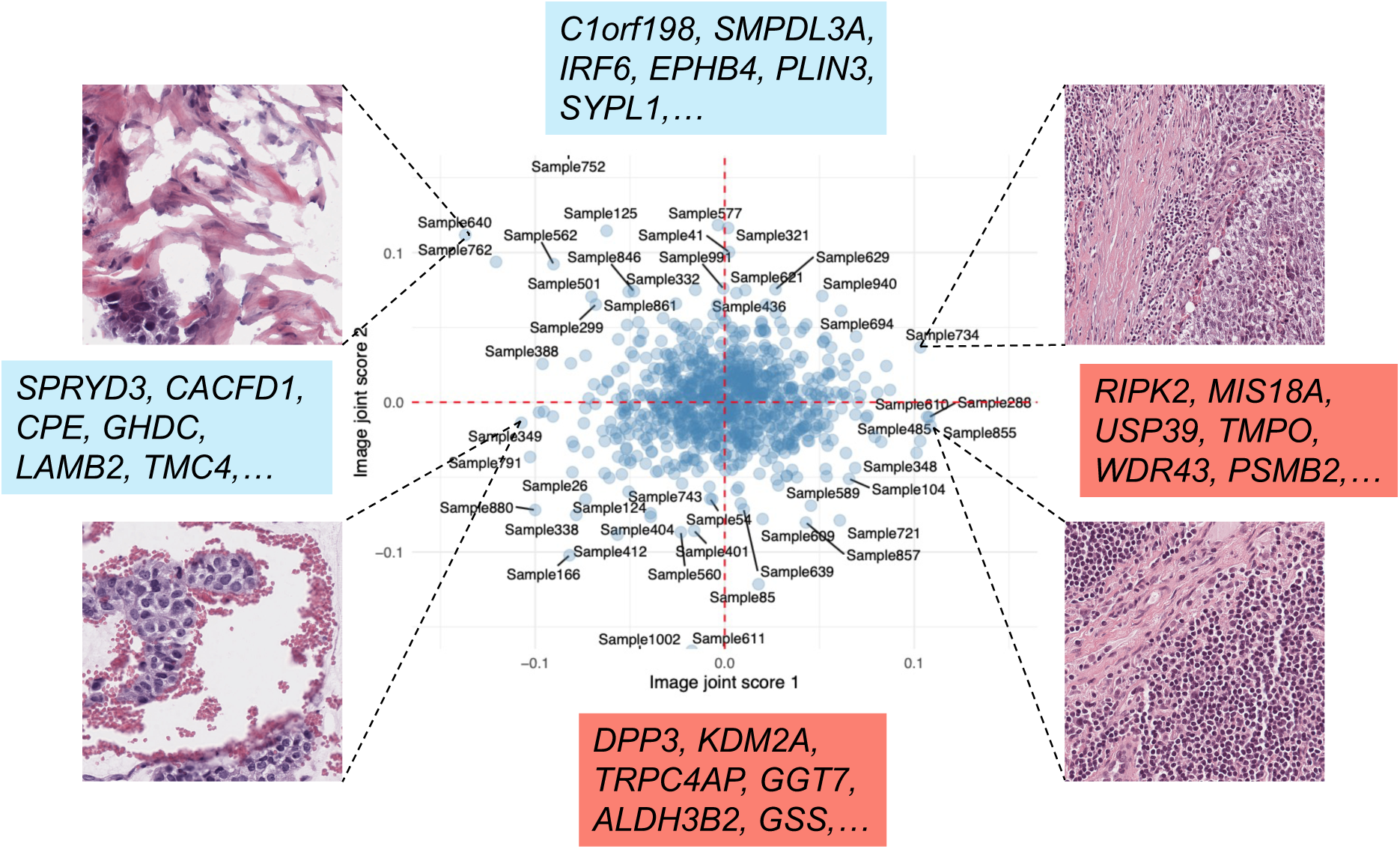
Joint latent structure in H&E image and gene expression identified by AJIVE in two-dimensional space defined by the image joint scores 1 and 2. Samples are plotted according to their joint scores with representative images from the positive and negative ends shown. Genes associated with the images are listed with either positive loadings (blue) or negative loadings (salmon). AJIVE identified joint components distinguishing densely packed small nuclei from enlarged heterogeneous nuclei (accompanied by substantial sparse empty regions), together with associated gene signatures.

We next investigated the associations between gene signatures and the corresponding image samples. Because of the negative correlation between the first joint score pair, image samples with positive scores were linked to genes with negative loadings, whereas image samples with negative scores were linked to genes with positive loadings. In the joint component 1, image samples with positive scores were associated with genes including *RIPK2*, *MIS18A*, *USP39*, *TMPO*, *WDR43*, and *PSMB2*, while image samples with negative scores were associated with genes including *SPRYD3*, *CACFD1*, *CPE*, *GHDC*, *LAMB2*, and *TMC4* (Figure 4). Notably, RIPK2 plays an important role in immune regulation, and increased RIPK2 activity has been reported in inflammatory breast cancer (IBC) [44]. Additionally, targeting RIPK2 with specific inhibitors has been proposed as a promising anti-tumor strategy [45].

To examine whether gene expression differed across subgroups defined by the joint component, samples were stratified based on their image joint score on component 1. The distribution of score values across all samples was approximately normal and centred around zero (Supplementary Figure S8A). We selected the top and bottom 10% of samples with the most extreme positive and negative scores to define two sample subgroups and compared their gene expression values. Interestingly, in the joint component 1, the expression of representative genes exhibited significant differences between the two subgroups. For example, the representative gene *RIPK2*, was highly expressed in the subgroup with a positive image joint score 1, whereas *SPRYD3*, was highly expressed in the subgroup with a negative image joint score 1 (Supplementary Figure S6).

On the joint component 2, image samples with a negative score displayed glandular-like structures with relatively large, irregular nuclei and visible nucleoli, whereas image samples with positive score showed nuclei arranged in sheet-like patterns with a greater presence of stromal tissue (Supplementary Figure S7). The second joint component exhibited a highly positive correlation between the pair of joint scores (Supplementary Figure S7). We therefore looked at the association between image samples with positive (negative) scores and genes with positive (negative) loadings. Positively scored image samples were linked with genes including *C1orf198*, *SMPDL3A*, *IRF6*, *EPHB4*, *PLIN3*, *SYPL1*, while negatively scored image samples were associated with genes including *GSS*, *ALDH3B2*, *GGT7*, *TRPC4AP*, *KDM2A*, *DPP3* (Supplementary Figure S7). Interestingly, IRF6 has been reported to be downregulated in highly invasive breast cancer cell lines, and its increased expression has been shown to suppress cell proliferation and tumorigenicity [46]. *KDM2A* has been implicated as an oncogene in breast cancer, and inhibition of KDM2A suppresses tumor growth [47, 48].

We conducted a similar subgroup sample analysis (Supplementary Figure S8B). Unlike what we observed for joint component 1, we observed less pronounced patterns. For example, the representative gene with a negative loading, *DPP3*, showed only modest differential expression (*p* = 0.22, *t* -test), whereas the representative gene with a positive loading, *C1orf198*, reached nominal statistical significance (*p* = 0.026, *t* -test). (Supplementary Figure S7).

We further examined the individual components identified by AJIVE, which are specific to each view and not shared across the two views. For image individual component 1, samples with extreme positive scores exhibited a relatively homogeneous background, with a higher proportion of eosin-stained (pink) stroma and sparse cellularity. In contrast, samples with extreme negative scores showed densely packed cells, marked variation in nuclear size, and disorganised tissue architecture (Supplementary Figure S9). We examined genes with extreme loadings in expression individual component 1. Genes with the top positive and negative loadings were predominantly enriched in RNA processing-related pathways, such as RNA splicing (Supplementary Figure S10).

## 6 Cross-method comparison

In this section, we examine the characteristics of the image patterns identified by each method, their associated gene signatures, the variance explained by each factor, and whether the methods recover the PAM50 subtype structure.

### 6.1 Comparison of visual patterns identified across integration methods

To gain insight into the information captured by different factors, we examined the image patterns identified by each factor across different methods (see representative images in Figure 5A).

**Figure 5.**
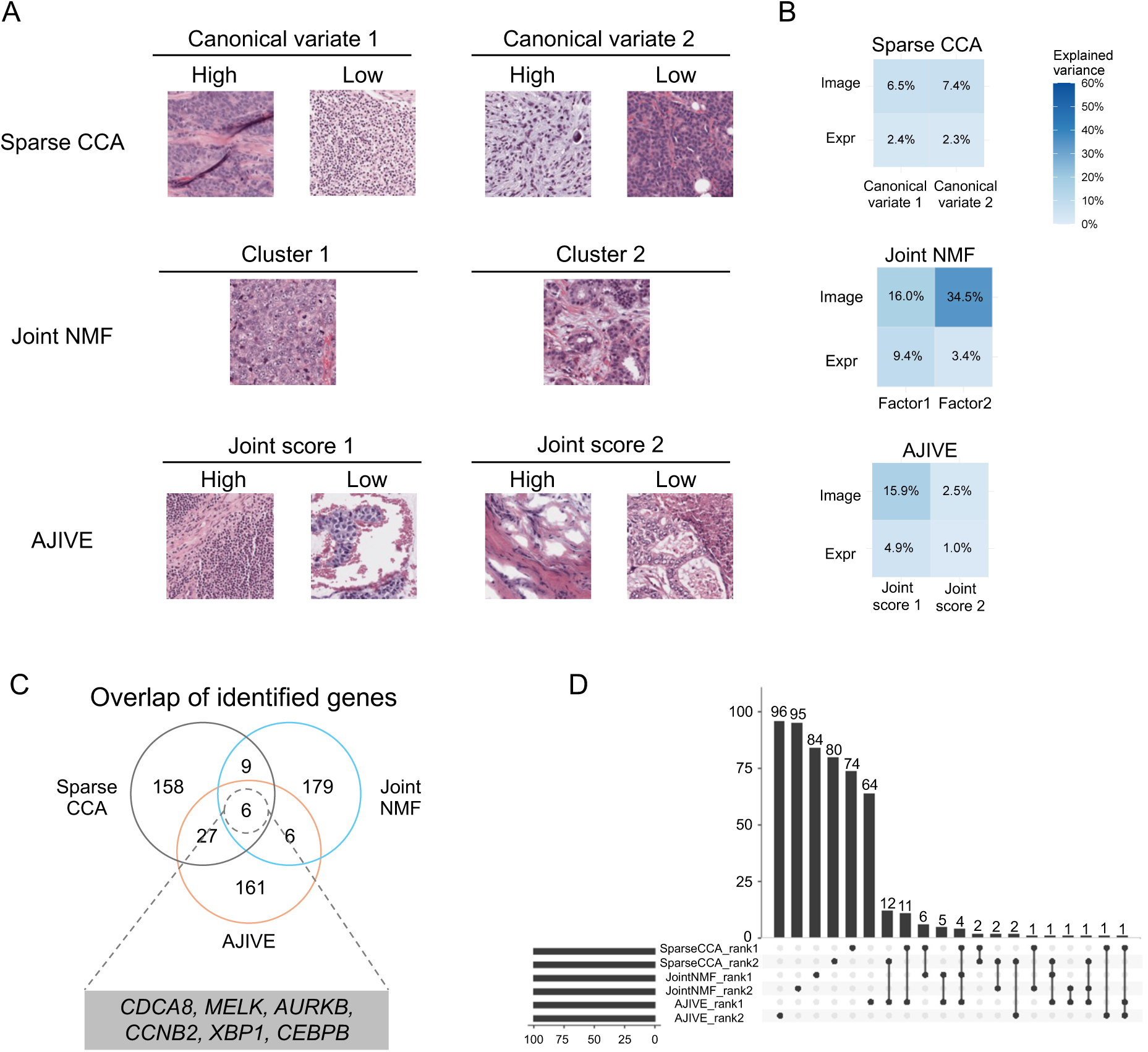
Comparison of integration results across methods. **A:** Representative images identified by each integration method are shown. For Sparse CCA: images with high and low values corresponding to each canonical variate; for Joint NMF: representative images for each cluster; and for AJIVE: images with high and low values associated with each joint score. **B:** Proportion of variance explained by each factor within each method. The rows represent the image and gene expression data, while the columns denote the factors identified by each method. **C:** Venn diagram showing the overlap of genes identified by the three methods, with six genes found in common, out of a selection of 100 genes per component. The common genes were identified as *CDCA*8, *MELK*, *AURKB*, *CCNB*2, *XBP* 1, *CEBPB*. **D:** Overlap of genes identified by each factor across the three method. The methods identified distinct representative image patterns and gene signatures, and captured different proportions of variance across components.

In sparse CCA, the first canonical factor clearly captured nuclear size: the image with high scores exhibited larger nuclei, whereas the image with low scores showed small, densely packed nuclei. For the second canonical factor, the image with high scores presented elongated, spindle-shaped cells with more dispersed nuclei and a pale purple background, indicative of stromal region. In contrast, the image with low scores displayed tightly arranged cells with high nuclear-to-cytoplasmic ratios and deeply stained nuclei (dark purple), with occasional adipocyte-like vacuoles in the background. (Figure 5A, Top row).

In Joint NMF, as noted earlier, the representative image from cluster 1 exhibited marked nuclear atypia, loss of glandular architecture, and minimal visible stroma. In comparison, the representative image from cluster 2 retained discernible glandular structures together with visible stromal tissue (Figure 5A, Middle row).

In AJIVE, the first joint component also captured characteristics in nuclear morphology. While this showed some similarity to Sparse CCA (both captured information in nuclear morphology), the overall captured patterns remained distinct. The image with high scores showed densely packed small nuclei with a minor presence of stromal tissue, whereas the image with low scores revealed enlarged and heterogeneous nuclei surrounded by small eosinophilic dots. For the second component, the image with high scores demonstrated sheet-like nuclear arrangements with an increased stromal presence, while the image with low scores exhibited glandular-like structures characterised by relatively large, irregular nuclei with visible nucleoli (Figure 5A, Bottom row).

We further compared the proportion of image variance captured by the latent factors across methods. Overall, Joint NMF captured a substantial proportion of the variance with its two factors (16.0% and 34.5% for Factors 1 and 2, respectively). While AJIVE captured a moderate amount of variance with its first joint component (15.9%), the second component contributed comparatively little (2.5%). In contrast, Sparse CCA distributed the small explained variance more evenly across canonical variates (6.5% and 7.4% for canonical variates 1 and 2, respectively) (Figure 5B).

### 6.2 Comparison of identified gene signatures

Next, we investigated the corresponding patterns in the gene expression data. We first assessed the overlap among genes selected by each method. Restricting each component to its top 100 genes ranked by loading magnitude, only six genes were shared across all three methods, including *CDCA8*, *MELK*, *AURKB*, *CCNB2*, *XBP1*, *CEBPB* (Figure 5C). These genes are primarily involved in cell proliferation and transcriptional regulation [49–54]. When restricting each component to its top 50 genes, only one gene, *AURKB* overlapped across all methods (Supplementary Figure S11A).

We further examined the genes associated with each factor across the three methods. A substantial proportion of genes were uniquely identified by individual factors, with only a small number shared across different factors (Figure 5D; Supplementary Figure S11B). To characterise the functional relevance of these top genes, GO enrichment analysis was performed on the top 100 genes from each factor. Interestingly, the genes associated with the first component of all three methods were consistently enriched in cell proliferation related pathways (Supplementary Figure S12).

Similarly, we quantified the proportion of variance in gene expression data explained by the corresponding latent factors. Overall, Joint NMF accounted for the relative larger proportion of variance (9.4% and 3.4% for Factors 1 and 2, respectively). In contrast, Sparse CCA and AJIVE explained comparatively smaller proportions of variance (Figure 5B).

### 6.3 PAM50-associated structures captured by Sparse CCA and Joint NMF

To further evaluate whether the low-dimensional representations captured biologically meaningful structure, we assigned PAM50 molecular subtypes to all samples based on their PAM50 gene expression. In the gene expression space derived from Sparse CCA and from Joint NMF, we observed discernible subtype-related structure. Specifically, the Basal subtype was clearly separated from the other groups (Figure 6A, B). In Sparse CCA, additional separation among non-Basal subtypes was also apparent compared to Joint NMF (Figure 6A, B). In contrast, the low-dimensional representation derived from AJIVE did not reveal obvious PAM50-associated structure (Supplementary Figure S13B, C). Interestingly, however, in the space spanned by the AJIVE expression individual components, some PAM50-related structure can be observed, with the Basal subtype showing separation from the other subtypes (Supplementary Figure S13D).

**Figure 6.**
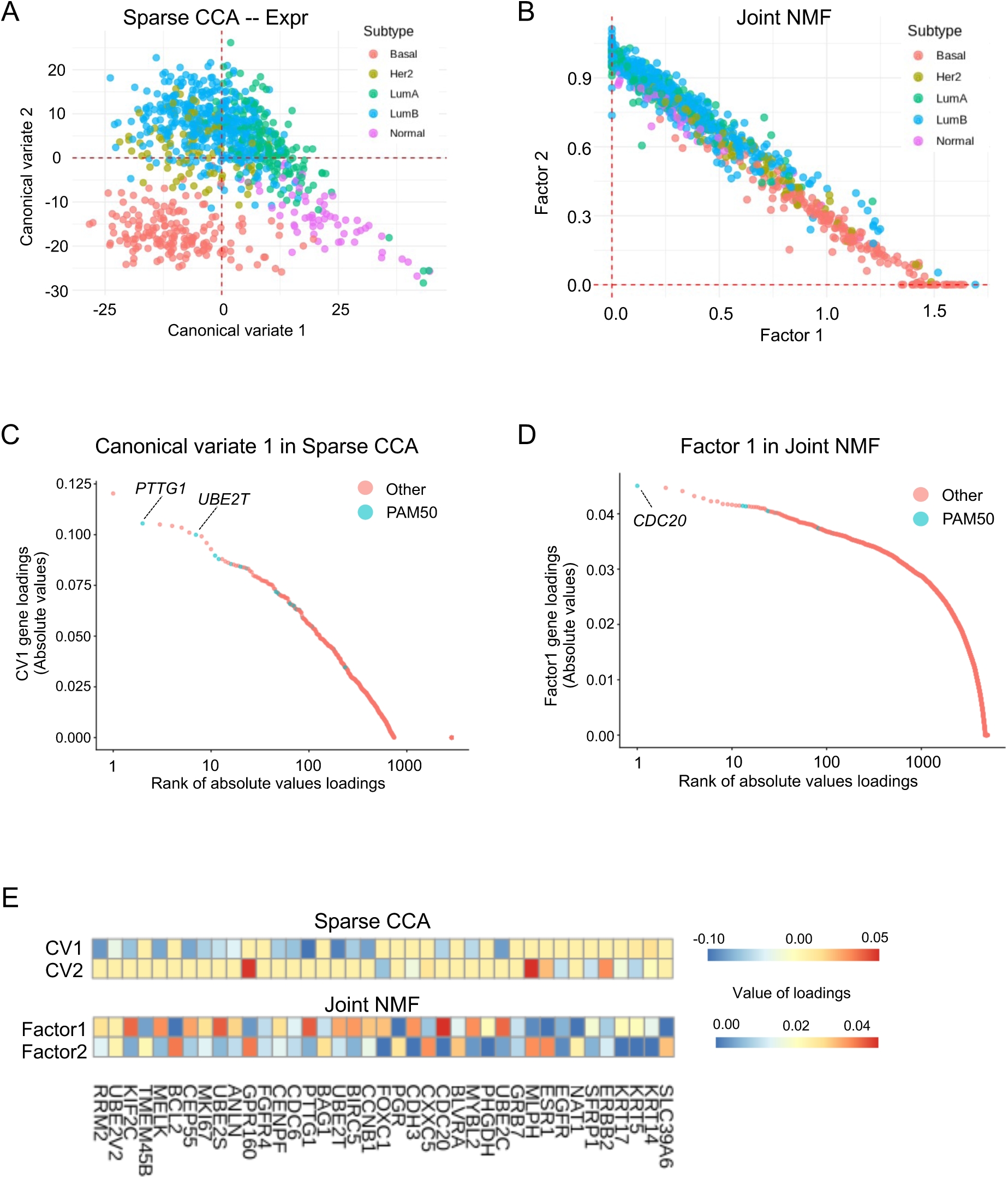
Sparse CCA and Joint NMF captured PAM50-associated structure in low-dimensional space. **A** and **B:** Sparse CCA canonical variates 1 and 2 corresponding to the gene expression view and Joint NMF Factors 1 and 2. Samples are coloured according to their PAM50 molecular subtypes. **C** and **D:** Gene loadings associated with either canonical variate 1 in Sparse CCA or Factor 1 in Joint NMF, ranked by absolute loading values. Genes are coloured according to PAM50 versus non-PAM50 membership. **E:** Heatmaps of PAM50 gene loadings associated with canonical variates 1 and 2 from Sparse CCA (top) and Factors 1 and 2 from Joint NMF (bottom). Sparse CCA and Joint NMF captured PAM50-associated latent structures, with different PAM50-related genes contributing to the corresponding latent spaces.

To investigate which genes are driving this subtype-related signal, we examined the top-loading genes associated with each factor. In the first factor of both Sparse CCA and Joint NMF, several PAM50 genes were present among the top 10 ranked loadings, including *PTTG1* and *UBE2T* in Sparse CCA, and *CDC20* in Joint NMF (Figure 6C, D). No PAM50 genes were observed among the top loadings of the second factor from Sparse CCA, Joint NMF, or either joint component from AJIVE (Supplementary Figure S14).

We further examined the loadings of all PAM50 genes associated with the two factors identified by Sparse CCA and Joint NMF. In Sparse CCA, the imposed sparsity constraint automatically selects a subset of genes with non-zero loadings, facilitating the identification of the most relevant features. The first canonical variate selected a higher number of PAM50 genes, with relatively large loading magnitudes for *PTTG1*, *UBE2T*, *RRM2*, *CCNB1*, and *UBE2C*. The second canonical variate involved fewer PAM50 genes, primarily including *GPR160*, *MLPH*, and *ERBB2*. For Joint NMF, two non-negative joint factors were obtained. The first joint factor was characterised by high loadings for genes such as *CDC20*, *KIF2C*, *UBE2S*, *PTTG1*, and *UBE2C*. The second joint factor showed larger loadings for *BCL2*, *GPR160*, *CXXC5*, *MLPH*, and *ESR1* (Figure 6E).

## 7 Discussion

In this review, we analysed the TCGA-BRCA datasets as a representative case study to compare three multivariate integration methods for integrating H&E images and gene expression data. While the specific biological findings reported here are derived from the TCGA-BRCA cohort only, the methodological differences observed among the three approaches extend more broadly to imaging-omics integration studies. The most notable finding of this review is that different methods captured different aspects of the underlying variation, providing complementary insights into the relationship between tissue morphology and molecular profiles. This can be attributed to differences in the underlying principles of each method, including modelling assumptions and optimisation strategies. Interestingly, PAM50 subtype structure was partially recovered within the latent spaces identified by Sparse CCA and Joint NMF, suggesting that biologically relevant subtype signals were embedded within the shared image-gene expression structure.

Multivariate integration methods offer an interpretable approach for linking morphological characteristics with gene expression profiles. With respect to the interpretability of gene-image associations, Sparse CCA provided intuitive framework by linking genes (through positive and negative loadings) to distinct subgroups of image samples. The sparsity constraint on the loading vectors facilitates the automatic selection of a subset of features, which further improves interpretability. Similarly, AJIVE enabled the association of genes with specific image samples based on gene loadings and image scores. In contrast, Joint NMF constraints both scores and loadings to be nonnegative. This nonnegativity leads to a parts-based representation, in which each factor captures additive components of the data. The interpretation of each factor can be informed by the set of genes associated with that factor.

Despite their utility, each method has methodological limitations that should be considered when interpreting the results. One limitation of Sparse CCA is that the sparsity constraint relaxes the orthogonality property of canonical variates, resulting in correlated latent dimensions. Similarly, the latent factors identified by Joint NMF are not constrained to be orthogonal and may therefore capture partially overlapping signals. Another limitation of NMF is its sensitivity to initialisation due to the non-convex nature of the optimisation problem. Although stable variants such as consensus NMF (cNMF) [55] exist for single-view settings, this issue can still persist in joint multi-view settings. Developing more stable joint NMF frameworks remains an important direction for future research. Moreover, neither Joint NMF nor AJIVE inherently perform feature selection, which may complicate biological interpretation when large numbers of genes contribute to a component. Incorporating sparsity constraints into these frameworks may enable selection of the most informative genes, thereby enhancing biological interpretability. However, sparsity and orthogonality are not always simultaneously achievable [12, 56]. Balancing these two properties remains an important direction for future methodological research.

Several limitations of this study should be acknowledged. First, our analyses were conducted at the bulk level. Image features were aggregated across multiple image patches, which may obscure intra-tissue morphological heterogeneity. Similarly, gene expression measurements represented average expression levels within the tissue. In addition, although paired H&E images and gene expression profiles were obtained from the same patient, they do not necessarily originate from the same tissue region. These factors may reduce the strength of image-gene associations, which could partly explain the relatively low correlations observed between paired canonical variates in Sparse CCA (Supplementary Figure S2A and S2B). Therefore, extending multivariate integration methods to spatially resolved transcriptomics is a promising direction for future research, as it enables direct mapping between spatial gene expression and corresponding image features, and may yield stronger and more precise gene-image associations.

## Resource availability

### Data availability

All datasets used in this study are publicly available from The Cancer Genome Atlas (TCGA-BRCA) via the GDC Data Portal (https://portal.gdc.cancer.gov/).

### Code availability

Preprocessed data and analysis scripts for reproducing the results in this study are available on GitHub (https://github.com/ChengyiMA/Multivariate-integration-of-image-omics).

## Declarations

### Author contributions

C.M. conceived the study, performed the formal analyses, and wrote and edited the manuscript under the supervision of K.-A.L.C. and J.M.. K.-A.L.C. contributed to the study conception and manuscript editing. J.M. edited the manuscript.

## Acknowledgements

We would like to thank Prof. J. S. Marron (University of North Carolina at Chapel Hill) for helpful discussions.

## Supplemental information

1. Document S1. Supplementary Methods and Supplementary Figures S1–S14.

