## Supplemental material for "Multivariate integration of histological images and gene expression data: a comparative review"

### S1 Supplementary Methods

Implementation details of multivariate integration methods For sparse canonical correlation analysis (Sparse CCA), centred and scaled gene expression and image feature matrices were used as input. Sparse CCA was implemented using the PMA R package (v1.2.4) ([1]). The default sparsity penalty parameters were adopted. The number of canonical variates was set to two to maintain consistency across all integration methods.

For joint non-negative matrix factorisation (Joint NMF), L2 transformed gene expression and image feature matrices were used as input. The number of latent factors was set to two, as this configuration yielded the highest silhouette score and the most stable sample clustering. To mitigate convergence to local minima, the factorisation was repeated 50 times with random initialisations, and the solution with the lowest objective function value was retained. Following Zhang et al. [2], the original MATLAB implementation was re-implemented in R. Samples were clustered based on the joint score matrix using $K$-means clustering. The number of clusters was determined by maximising the silhouette score across a range of candidate values of $K$. The optimal number of clusters was then used for downstream analysis.

For angle-based joint and individual variation explained (AJIVE) [3], centred and scaled gene expression and image feature matrices were used as input. Two joint components were identified. The Python implementation of AJIVE (v0.2.0) was employed for the analysis.

Input data were tailored to satisfy the modelling assumptions of each method. Sparse CCA and AJIVE require centred and scaled inputs to ensure comparability across variables, whereas Joint NMF imposes non-negativity constraints on the input data and therefore was applied to L2-normalised non-negative matrices.


### S2 Supplementary Figures

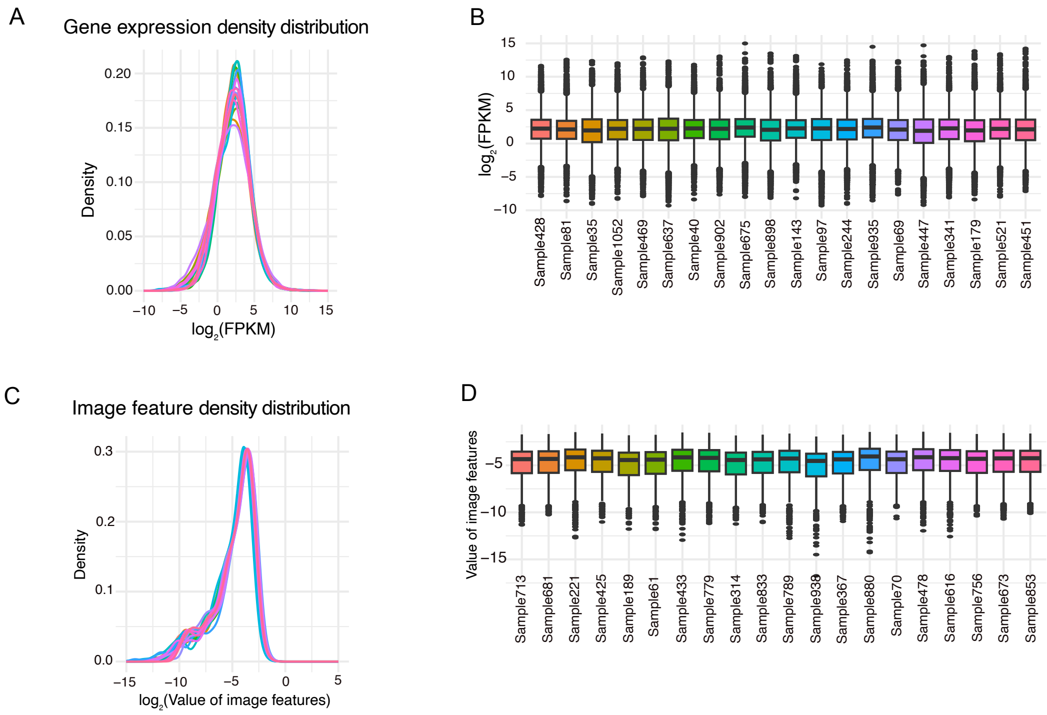


Figure S1: Feature distributions across samples after preprocessing. (A) and (B) Gene expression distributions shown as density plots and boxplots across 20 samples. (C) and (D) Image feature distributions shown as density plots and boxplots across 20 samples. *Samples showed consistent feature distributions after preprocessing.*

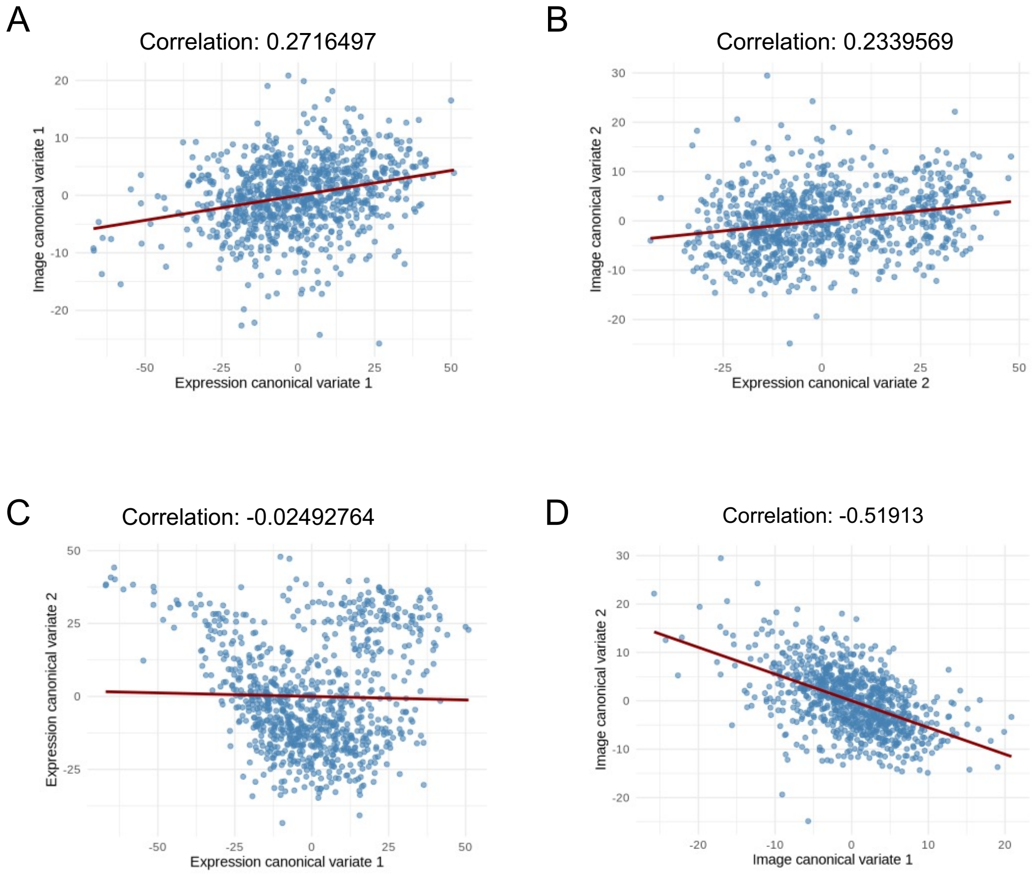


Figure S2: Correlations between image and expression canonical variates. (A) and (B) Correlations between paired image and expression canonical variates 1 and 2, respectively. (C) Correlation between expression canonical variates 1 and 2. (D) Correlation between image canonical variates 1 and 2. *Paired image and expression canonical variates showed positive correlations, while correlations were also observed between image canonical variates 1 and 2.*

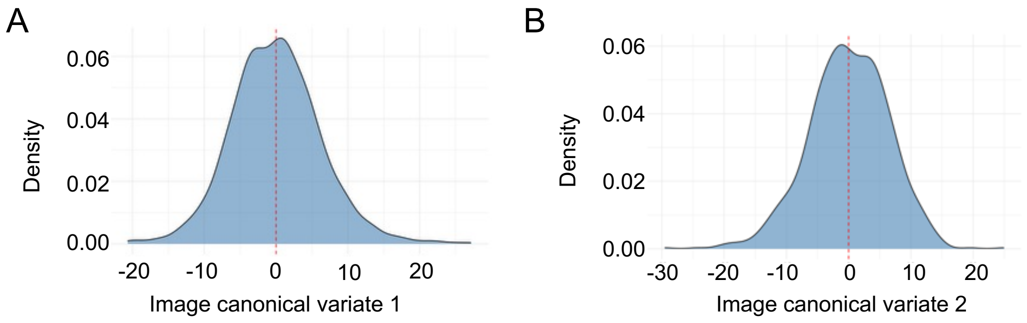


Figure S3: Sample distributions along image canonical variates 1 and 2. (A) and (B) Density distributions for canonical variates 1 and 2, respectively. *Sample distributions along the image canonical variates were approximately symmetric and centred at zero.*

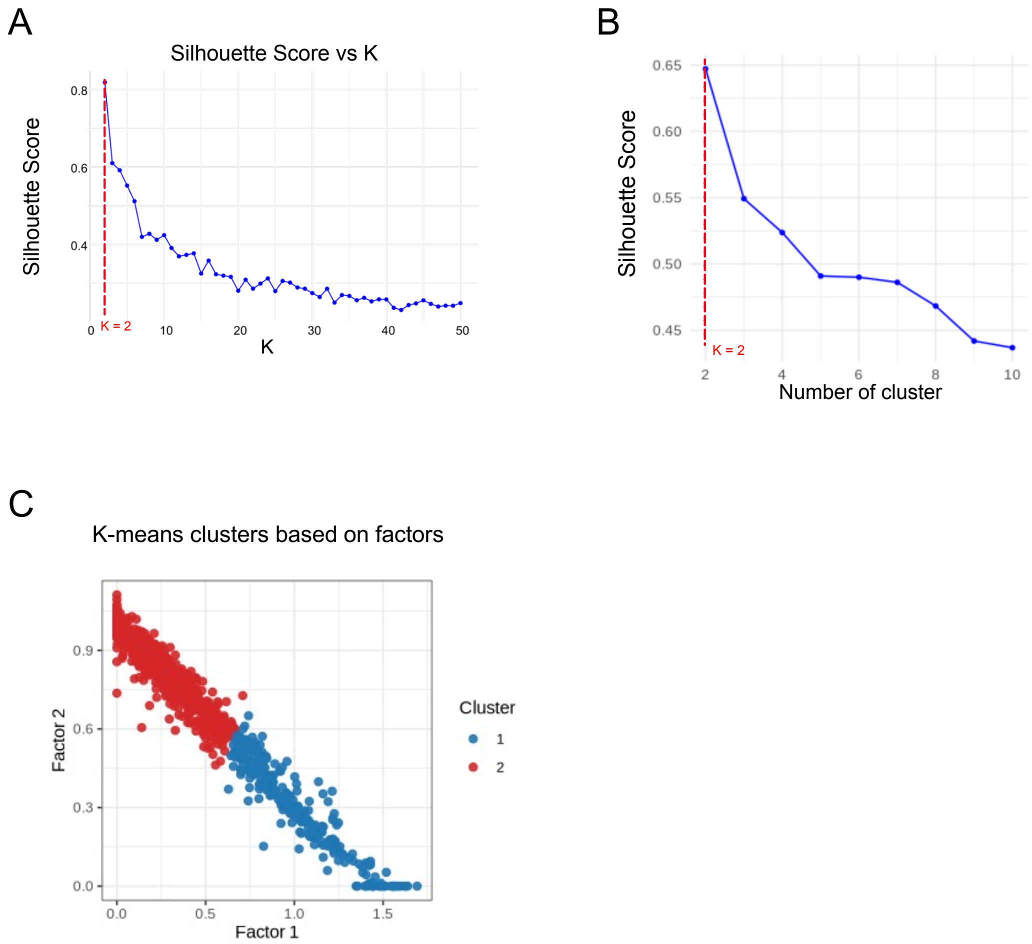


Figure S4: Factor selection and clustering in Joint NMF. (A) Silhouette scores for selecting the number of factors (K), with K=2 yielding the highest score. (B) Silhouette scores for k-means clustering of joint scores across different numbers of clusters, with two clusters performing best. (C) Scatter plot of samples in the Joint NMF factor space (Factor 1 vs Factor 2), coloured by k-means cluster labels. *k-means clustering on the joint score matrix (from Joint NMF) identified two clusters.*

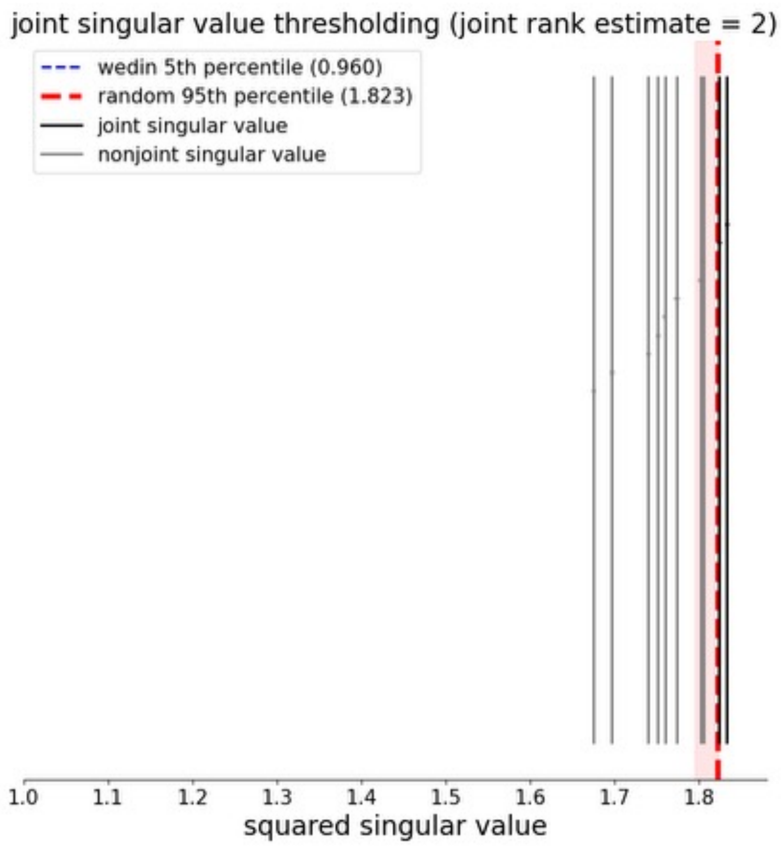


Figure S5: AJIVE joint rank estimate, showing a joint rank of 2.

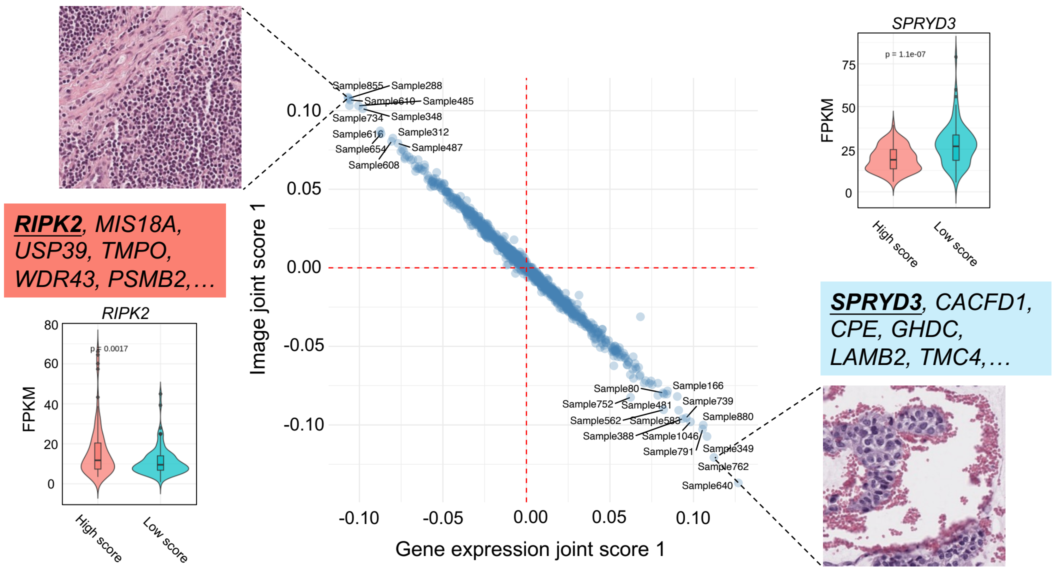


Figure S6: Joint component 1 identified by AJIVE. Samples were projected into a two-dimensional space defined by the first paired joint scores. In this component, gene expression and image joint scores were completely negatively correlated. Representative images from the positive and negative ends were shown. Genes associated with the images were listed with either positive loadings (blue) or negative loadings (salmon). The representative genes *RIPK2*, *SPRYD3* showed differential expression between the two subgroups defined by the image joint component 1. *The image joint component 1 captured nuclear morphology and associated gene signatures.*

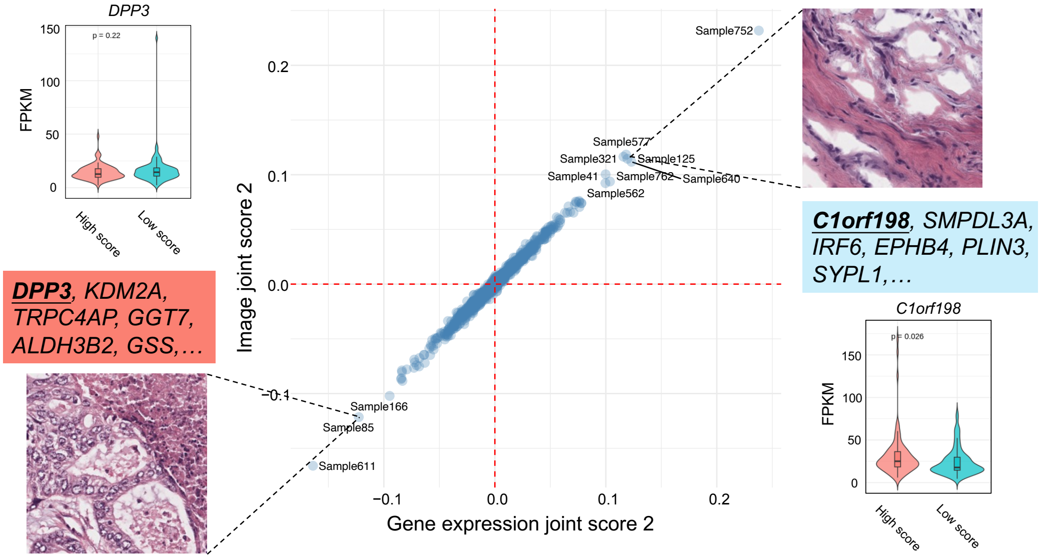


Figure S7: Joint component 2 identified by AJIVE. Samples were projected into a two-dimensional space defined by the second paired joint scores. Gene expression and image joint scores were completely positively correlated in this component. Representative images from the positive and negative ends were shown. Genes associated with the images were listed with positive loadings (blue) or negative loadings (salmon). The representative gene *DPP3* did not show significant differential expression between the two subgroups defined by image joint component 2, whereas *C1orf198* only marginally reached statistical significance. *The image joint component 2 captured sheet-like nuclear arrangements from enlarged irregular nuclei with visible nucleoli, together with associated gene signatures.*

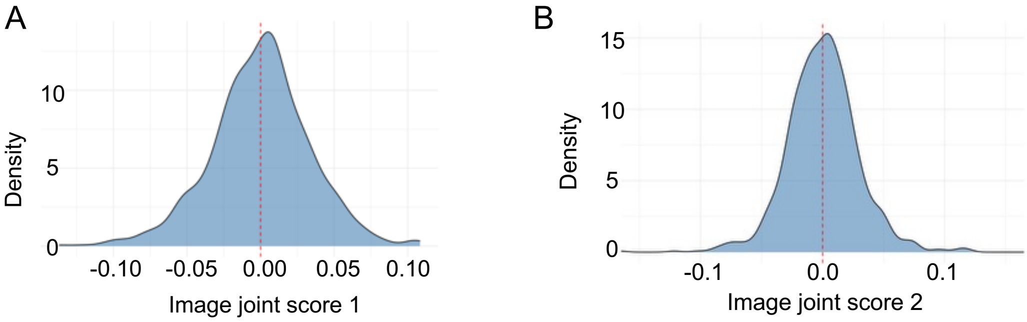


Figure S8: Sample distributions along image joint components 1 and 2 from AJIVE. (A) and (B) Density distributions of samples along image joint components 1 and 2, respectively. *Sample distributions along the image joint components were approximately symmetric and centred around zero.*

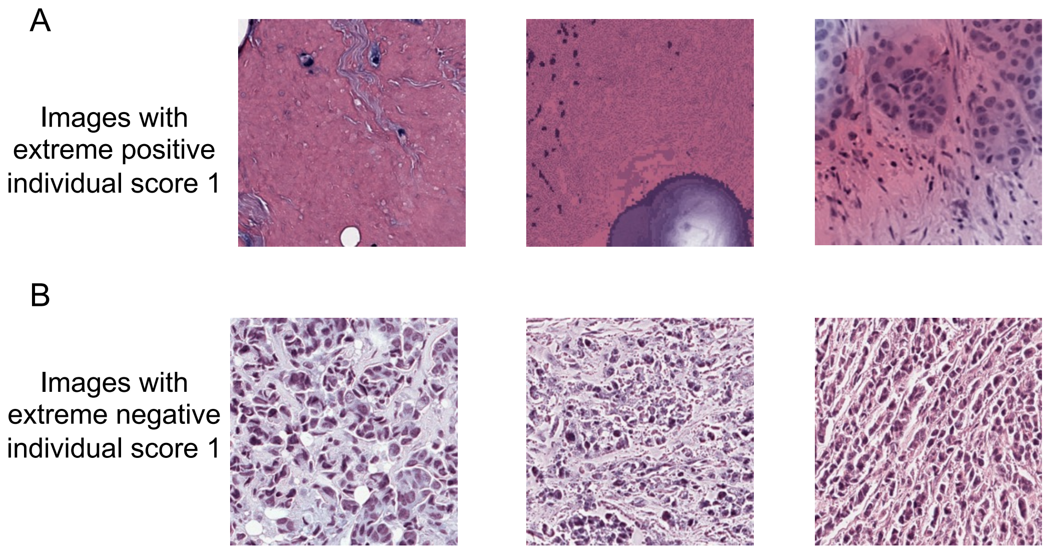


Figure S9: Image individual structures identified by AJIVE. (A) and (B) Representative images with extreme positive and negative values of image individual score 1, respectively. *Image individual components captured image-specific morphological structures which were not shared with gene expression data.*

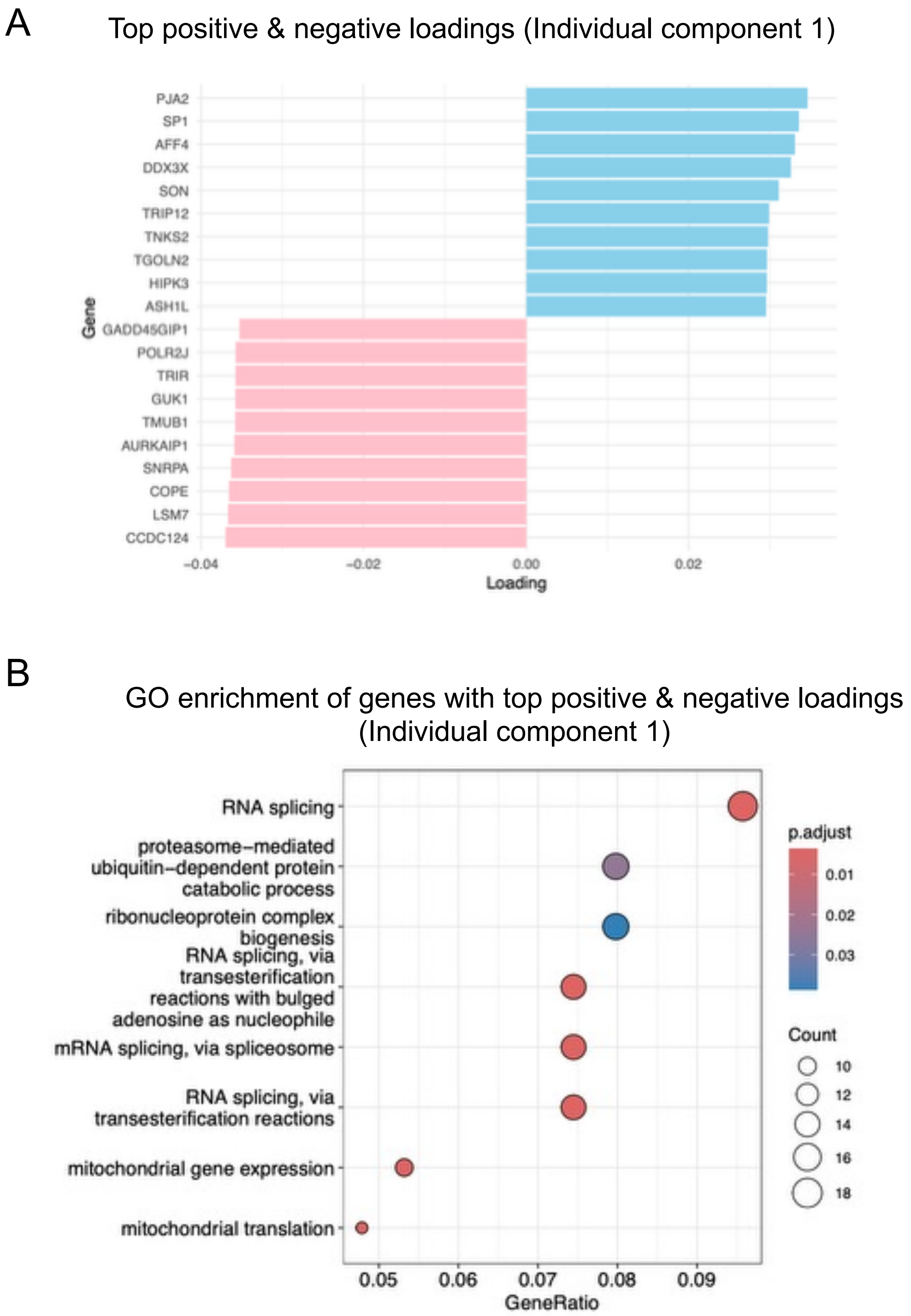


Figure S10: Individual structures of gene expression data identified by AJIVE. (A) Genes with extreme positive and negative loadings on gene expression individual component 1. (B) Go enrichment of genes with top positive and negative loadings associated with (gene expression) individual component 1. The top 100 genes were selected for positive and negative loadings, respectively. *Gene expression individual components captured gene expression variation not shared with image data, with associated genes primarily enriched for RNA splicing-related pathways.*

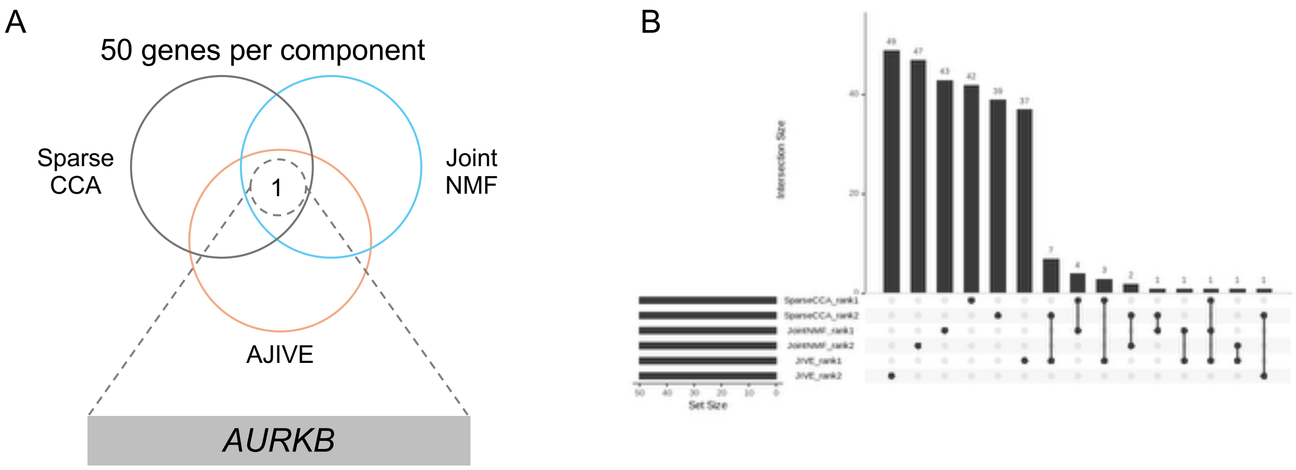


Figure S11: Common genes identified by each method (50 genes per component). (A) Venn diagram showing gene overlap across the three methods, with *AURKB* identified as the only common gene. (B) UpSet plot showing gene overlap across components and methods. *Using 50 genes per component, the three methods identified largely distinct gene sets, with only one overlapping gene (AURKB) across methods.*

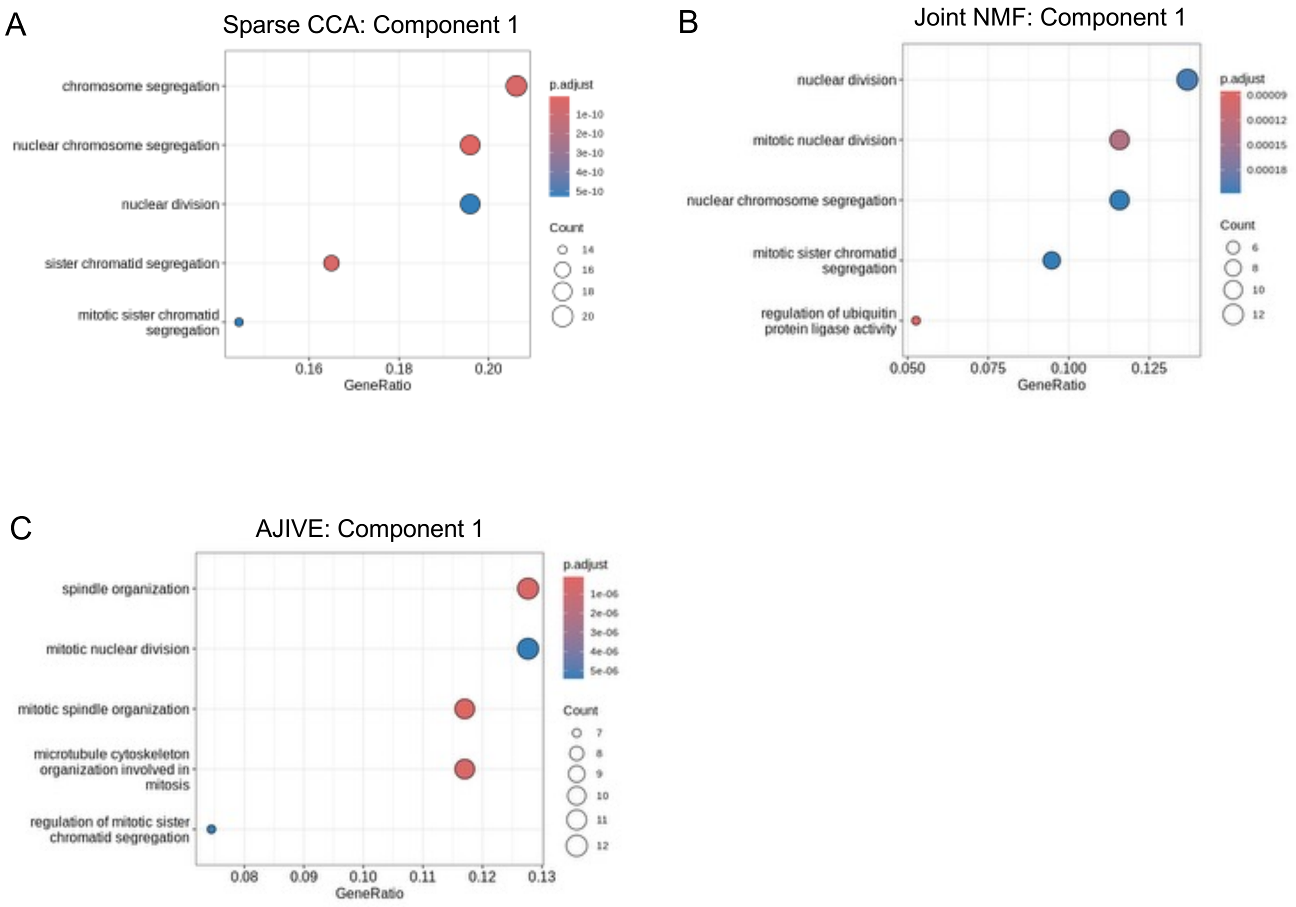


Figure S12: Enriched GO terms for genes selected in each component (100 genes per component). (A) Sparse CCA Component 1; (B) Joint NMF Component 1; (C) AJIVE Component 1. No GO terms were enriched for Component 2 across Sparse CCA, Joint NMF, or AJIVE. *Across methods, GO enrichment of genes selected in Component 1 (100 genes per component) consistently highlighted nuclear division-related pathways, while no enrichment was observed for Component 2.*

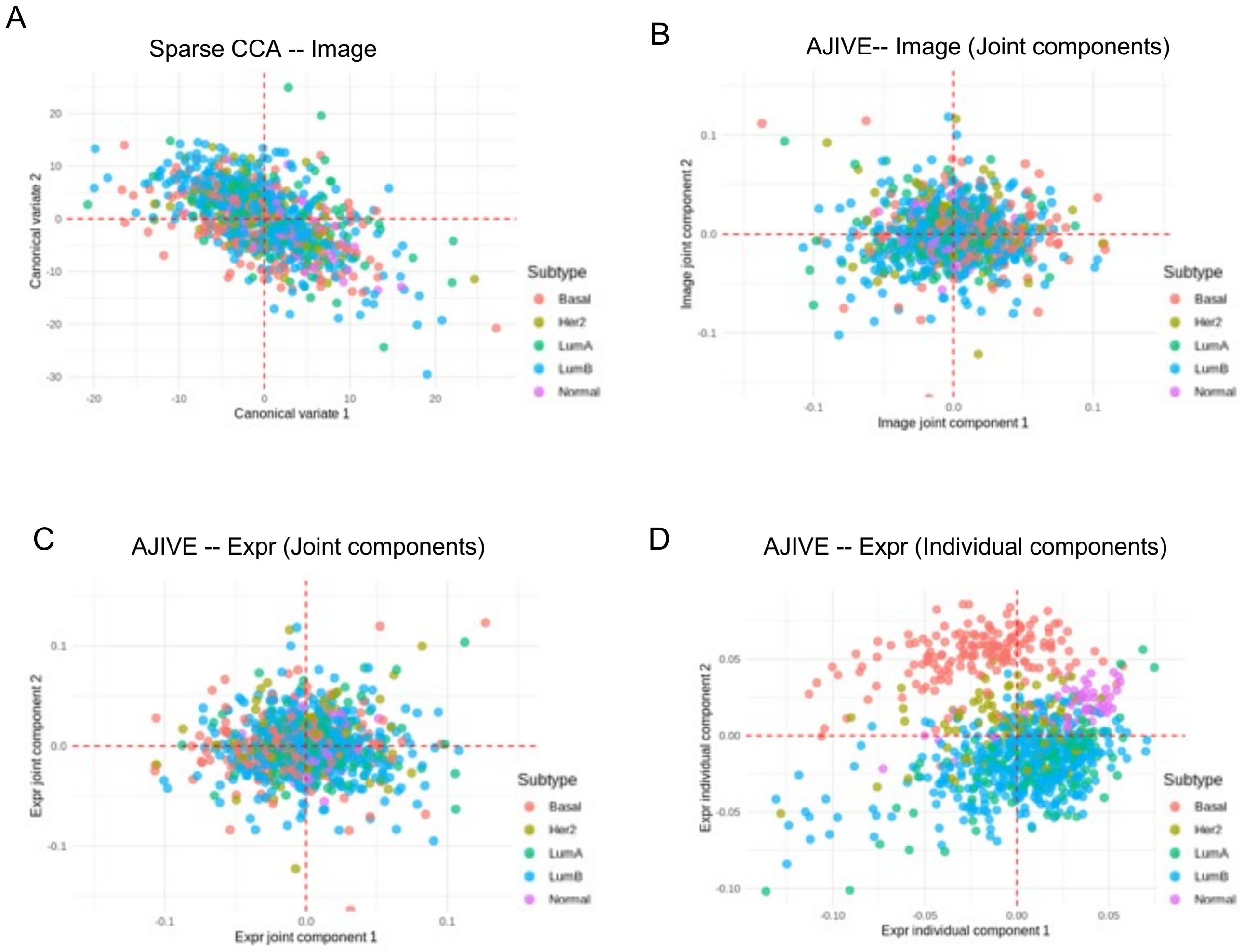


Figure S13: Sample scatter plots coloured by PAM50 molecular subtypes. (A) Sparse CCA image space (Canonical variates 1–2). (B) AJIVE image joint space (Components 1–2). (C) AJIVE gene expression joint space (Components 1–2). (D) AJIVE gene expression individual space (Components 1–2). *The AJIVE gene expression individual space captured PAM50-associated structure.*

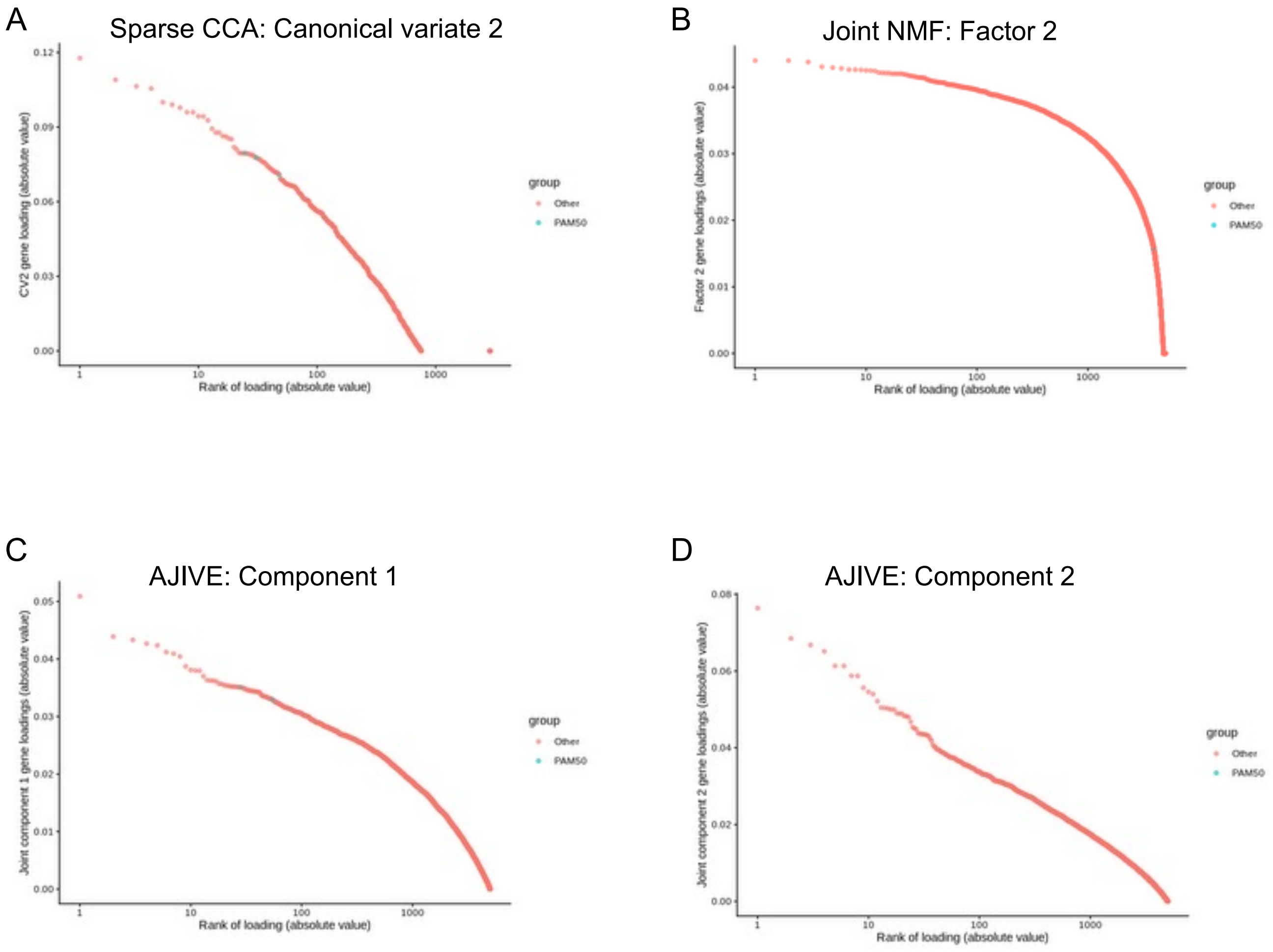


Figure S14: Gene loadings ranked by absolute values. (A) Sparse CCA, Canonical variate 2; (B) Joint NMF, Factor 2; (C) AJIVE, Component 1; (D) AJIVE, Component 2. Genes were coloured by PAM50 versus non-PAM50 membership. *Across these components, the top-ranked genes did not show strong enrichment for PAM50-related genes.*
